# Energy quality shapes biodiversity across coastal oceans

**DOI:** 10.64898/2026.09.23.753818

**Authors:** Simon J. Brandl, Nicole R. Foster, Robert F. Semmler, Ingrid Vasconcellos Bunholi, J. Emmett Duffy, Zhanfei Liu, Sean R. Connolly, Matthieu Leray, Jonathan S. Lefcheck, Katelyn DiBenedetto, Emily R. Anderson, David M. Baker, Ximena Boza, João Canning-Clode, Valentina Cardona, Rachel Collin, Lara Denis-Roy, Graham J. Edgar, Sonia K.M. Gueroun, Isis Guibert, Dean S. Janiak, Christopher P. Meyer, Matthew B. Ogburn, Angeleen M. Olson, Luciano Pastorelli, Christopher J. Patrick, Carolyn Prentice, Michael Rappé, Yoshimi M. Rii, Mathew Seymour, Ximena Velez-Zuazo, Jianhong Xue, Jordan M. Casey

## Abstract

Earth’s biodiversity is distributed unevenly, typically peaking in warm, vegetated, and stable environments. This has frequently been linked to energy availability, which boosts productivity, bolsters populations, and facilitates coexistence. Here, we examine the relationship between energy and biodiversity across coastal oceans using particulate organic matter biogeochemistry and environmental DNA. We reveal that energy quantity (concentrations of carbon, nitrogen, and hydrolyzable amino acids) and salinity, two variables that co-vary with freshwater inflow, jointly predict species richness at regional and global scales. Biodiversity was lowest at sites and locations with high energy inflow and low salinity, suggesting that environmental filtering associated with the osmotic stress induced by freshwater inflow outweighs raw resource availability in governing species richness. Independent of this gradient, however, energy quality (defined via C:N ratios, δ^13^C values, and amino acid profiles) was associated with higher biodiversity. Labile, protein-rich, marine-derived resources supported higher biodiversity across all taxa at the regional scale, and a larger number of planktonic consumer taxa at both regional and global scales. High energy quality sites were also enriched in key planktonic groups such as calanoid and cyclopoid copepods, suggesting that these sites act as hotspots of planktonic biodiversity across coastal seascapes. Our findings suggest that few species can directly harness the plentiful resources provided by terrestrial subsidies in coastal oceans because they arrive in low salinity waters. In turn, high-quality food attracts a diverse range of consumers, which may seed the patchy foraging hotspots that characterize open-water food webs. Altered coastal hydrodynamics and biogeochemistry may therefore affect nearshore biodiversity, food webs, and fisheries.

## Introduction

Life on our planet is sustained by electromagnetic energy radiating from the sun, which enables the biochemical synthesis of organic matter. Thus, the quantity of available energy, be it electromagnetic or chemical, is a crucial determinant of plant and animal communities across scales (1), including the paradigmatic decline of biodiversity towards Earth’s cold and seasonally variable poles (2, 3). Indeed, Hutchinson (4) suggested that fewer species persist in high latitudes due to limited solar irradiance and, as a result, lower primary producer biomass, a lack of structure-forming vegetation, and little food for consumers. This pattern has been confirmed across taxa, continents, and biomes, ranging from major groups of terrestrial plants and vertebrates (5, 6) to arthropods (7, 8) and marine invertebrates (9, 10). Despite continued debate regarding the shape, scale-dependency, consistency, and underlying mechanisms of the ‘species-energy relationship’ (1, 11), as well as its relative contribution to global biodiversity patterns compared to geological legacies and evolutionary forces (12), there is broad scientific support for its basic tenet: systems with more available resources, whether measured as irradiance, productivity, landscape greenness, or standing biomass, generally support more species (2, 13).

More than 70% of Earth’s surface is covered by ocean, which presents markedly different circumstances for species-energy relationships and their mechanisms compared to terrestrial environments (14). First, geological and biogeographical drivers of biodiversity patterns tend to have a smaller footprint in the ocean compared to land (but see (15)), as glacial fluctuations or tectonic events are less likely to affect biota with high dispersal potential due to pelagic life stages (12). Second, while low solar irradiance and temperature clearly contribute to the decline of biodiversity around the poles (resulting in apparent species-energy relationships) (16), marine primary production is strongly regulated by ocean circulation and nutrient supply (17). These physical and chemical forces can obscure latitudinal trends in phytoplankton diversity outside polar regions (18–20). A similar pattern is visible for zooplankton, where sharp declines in the number of species at the highest latitudes contrast with the relatively even distribution of species richness across temperate and tropical clines, suggesting factors such as salinity, food availability, and biotic interactions as important drivers of diversity (21–23). Despite the similarity in global producer and consumer patterns, however, the relationship between proxies of chemical energy availability (namely chlorophyll *a* concentrations or phytoplankton biomass) and zooplankton diversity is surprisingly weak (16, 22), raising questions about whether the quantity of available chemical energy supports higher consumer diversity as observed in many terrestrial systems (6, 24). This may, in part, be tied to the third hallmark of open-water marine systems: in contrast to most terrestrial and some benthic systems (e.g., kelp forests), primary producers do not usually fill a dual functional role as food producers and habitat builders. Phytoplankton provide sustenance, but no structural framework for animals to inhabit.

Relationships between energy availability and consumer communities in marine open-water systems are relevant for understanding energy fluxes that underpin the world’s fisheries (25). Predators in both coastal and pelagic open-water systems generally exist in a resource-poor environment (26), but they reliably converge on spatially variable and temporally transitory patches of high resource concentrations where a wide range of small, planktonic consumer species sustain many large bodied animals (27–29). Where, when, and how these ephemeral hotspots of biological activity emerge has remained enigmatic, despite their vital importance for life in marine systems and the services they provide to billions of people (26). Physical forcing (through convergent or divergent fronts) generally results in biogeochemical gradients that may affect consumer biodiversity and ultimately promote marine foraging hotspots. This is the case at the global scale, where variation in ocean circulation patterns and resulting nutrient fluxes (e.g., nutrient upwelling) shape marine biodiversity and foster the synthesis of chemical energy by phytoplankton to fuel pelagic productivity (30–32). However, proxies of energy availability (most commonly chlorophyll *a* concentrations) frequently fail to predictably correlate with biodiversity or biological activity in marine systems (27, 33, 34), and at finer spatial scales, relatively little is known about how foraging hotspots form.

In both contexts (i.e., species-energy theory and resource patchiness in open water systems), one factor that has received relatively little attention is the nutritional quality and bioavailability of resources. Resources typically fall on a spectrum from ‘fresh’ labile food that is protein- and micronutrient-rich and readily assimilated by a wide range of organisms to ‘old’ recalcitrant, carbon-rich material that can only be re-mineralized by specialized organisms (35, 36). These attributes are not well captured by standard methods used to assess resource pools in marine systems (e.g., remote sensing of chlorophyll *a* concentrations, bulk organic carbon measurements, field measurements of turbidity in marine environments). Yet, as indicators of “resource quality” (37, 38), they may be important in structuring consumer communities and their functioning as suggested by the principles of ecological stoichiometry (39). The conceptual and mechanistic distinction between bulk standing resources (hereafter termed “energy quantity”) and their origin, bioavailability, and nutritional value (hereafter termed “energy quality”, which captures the stoichiometric value of resources) may be especially relevant in surface waters of dynamic coastal ecosystems. In these systems, multiple sources of carbon and nitrogen collide to create complex seascapes of varying energy quantity and quality with strong cascading effects through marine food webs (40). Thus, biodiversity in coastal open-water systems may be governed by how much bulk energy is available (the species-energy relationship), how nutritionally accessible or valuable the resources are (ecological stoichiometry), and where the most favorable combination of these properties occurs at the seascape scale (resource patchiness). However, the intersection of these three broad ecological principles has not been explored in marine ecosystems.

Here, we test the relationship between consumer biodiversity and the system-wide energy landscape using water samples collected across a global coastal ocean network. At 60 sites distributed across twelve coastal locations that span tropical and temperate regions between -42.6° (Tasmania, Australia) and 50.2° (British Columbia, Canada) latitude (Fig. 1A), we measured diversity using environmental DNA (eDNA) metabarcoding and quantified bulk organic matter (i.e., *energy quantity*) and its availability and nutritional value (i.e., *energy quality*) using biogeochemical analyses of particulate organic matter. We then tested correlations between biodiversity and energy quantity and quality, alongside salinity and water temperature at regional and global scales and compared analyses using all detected taxa (i.e., taxonomically-uninformed biodiversity) and subsets of planktonic consumer taxa in open-water ecosystems.

**Figure 1.**
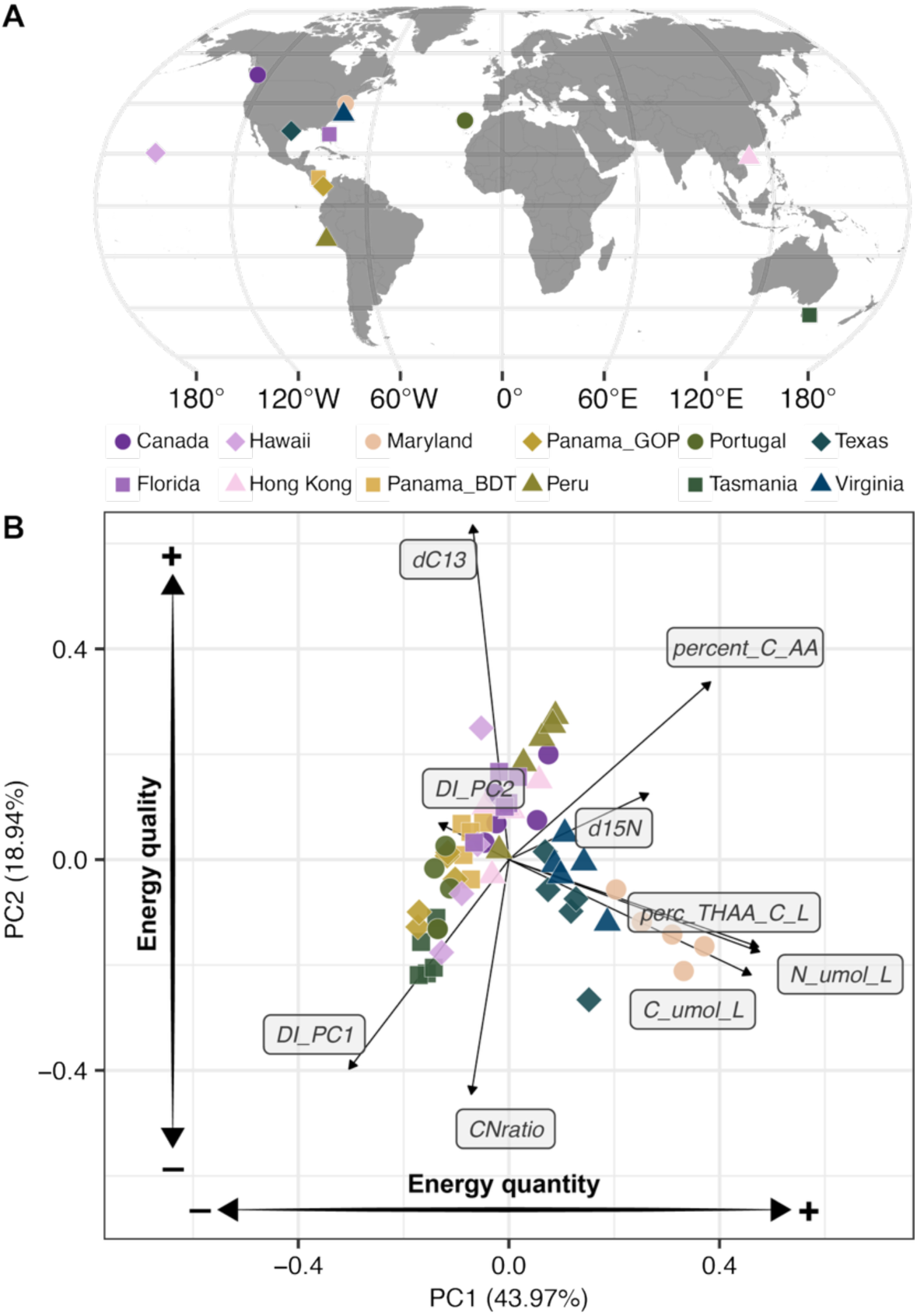
Distribution of sites and their biogeochemical profiles. (A) Map of the study locations, with points indicating each location (as the centroid of five sites within each location). (B) Principal Components Analysis (PCA) ordination of biogeochemical variables, showing differentiation of sites along the first two axes, which broadly align with gradients in energy quantity (PC1) and quality (PC2). Points represent average values from three replicate samples for each site (N = 5) within each location (N = 12), with locations being identified by unique combinations of shapes and colors (see legend). Panama_GOP = Gulf of Panamá, Panamá, Panama_BDT = Bocas del Toro, Panamá. Black lines and labels represent the vector loadings. *N_umol_L* = μmol N L^-1^; *C_umol_L* = μmol C L^-1^; *perc_THAA_C_L* = % total hydrolyzable amino acids C L^-1^; *d15N* = δ^15^N; *percent_C_AA* = % amino acids C L^-1^; *d13C* = δ^13^C; *CNratio* = C:N ratio; *DI_PC1* = degradation index axis 1; *DI_PC2* = degradation index axis 2. The degradation index was calculated based on the composition of total hydrolyzable amino acids (see Methods).

## Results

A principal component analysis (PCA) of particulate organic matter (POM) samples across sites and locations resolved two interpretable, orthogonal axes, which were characterized by relative concentrations of organic carbon (μmol C L^-1^), nitrogen (μmol N L^-1^), and total hydrolyzable amino acids (%THAA) in particulate matter samples (PC1), and C:N ratios, δ^13^C values, and an index of degradation based on amino acid profiles (41) (see Methods) on PC2. Given this, we defined *energy quantity* as a latent variable index denoted by PC1-scores, and *energy quality* as a latent variable index denoted by PC2 scores. While these synthetic indices do not directly quantify caloric potential or nutritional value in a metabolic sense, they represent informative proxies of the broader available resource environment. In this context, our energy quantity axis is comparable, in principle, to variables such as temperature, plant biomass, or NDVI, while our energy quality axis represents a rarely investigated, but potentially important descriptor of how bioavailable and nutritionally advantageous ambient energy is for consumers, providing a stoichiometric perspective.

We tested for the effects of energy quantity, energy quality, salinity, and water temperature on marine biodiversity, which was calculated based on incidence-frequency based estimates of amplicon sequence variants (ASVs) across five samples taken at each site (42, 43) at two different scales (across locations globally, and within each location). At the global scale (i.e., across our 12 locations), energy quantity was negatively associated with biodiversity (posterior parameter estimate *β* = −0.78 [-0.86 lower credible interval, −0.70 upper credible interval]; 100% probability of negative effect), while water temperature and energy quality had no clear effect (Fig. 2). Salinity, which was fit as a covariate in a separate model due to high collinearity with energy quantity (Fig. S1), showed a strong opposite correlation with biodiversity, with more saline locations harboring more species (0.72 [0.66, 0.78]; 100% probability). Globally, when approximating biodiversity through ASV richness across taxa, locations with more available energy in less saline waters support fewer species. Importantly, this pattern was not dependent on a discrete dichotomy between estuaries and marine sites: the same models run with a reduced dataset of locations with an average salinity of >30 again showed energy quantity (-0.57 [-0.74, −0.40]) and salinity (0.37 [0.20, 0.54]) as clear correlates of biodiversity.

**Figure 2.**
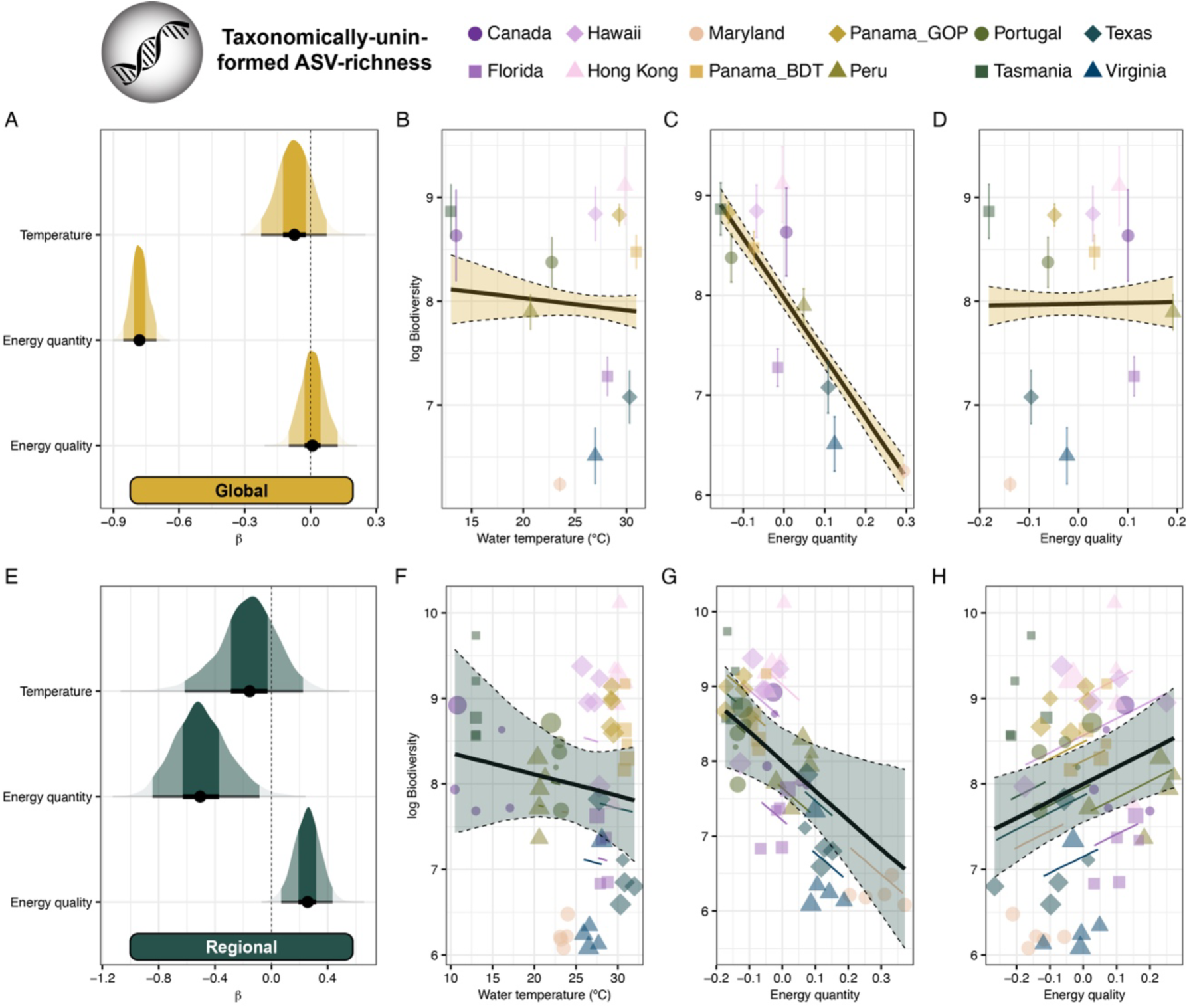
Relationships between diversity and environmental predictors across scales. (A and E) Posterior parameter distributions from Bayesian models indicating the strength and direction of scaled predictor effects in the global (A) and regional (E) models. All predictor variables were scaled and centered prior to the analysis for comparability. Shading and black caterpillar bars represent 50% and 95% credible intervals. (B – D and F – H) Partial effects plots of predictor variables. For the global model (B – D), points and error bars represent the weighted means and standard errors of the Chao2 estimator of amplicon sequence variant (ASV) richness, while the thick black line and dashed ribbon represent the predicted partial posterior fit derived from the model. For the regional model (F – H), points represent the weighted Chao2 estimator of ASV richness at each site as derived from the five replicate samples, with the size of the point reflecting the weight of each observation. The thick black line and dashed ribbon represent the predicted marginalized partial posterior fit derived from the model, while colored, solid lines represent the conditional slopes for each location. Combinations of shapes and colors correspond to the twelve locations (see legend). Panama_GOP = Gulf of Panamá, Panamá, Panama_BDT = Bocas del Toro, Panamá.

In contrast to the global analysis, we found clear and opposing effects of energy quantity and energy quality on biodiversity within regions. Specifically, energy quantity again showed a negative effect (-0.49 [-0.84; −0.08]; 99.9% probability) that was mirrored by an even stronger, opposing effect of salinity in a separate model (0.65 [0.34, 0.95]; 100% probability). In contrast, energy quality showed a clear, independent positive relationship with biodiversity at the regional scale (0.26 [0.07, 0.44]; 95.8% probability), while water temperature again had no effect. Local biodiversity evidently increases in more saline, oceanic waters, but it is independently boosted by the availability of labile, marine-derived, protein-rich organic matter (Fig. 2). This independent effect of energy quality on biodiversity at the regional scale was further bolstered by the model outcomes from the dataset with only marine salinities. In this dataset (N = 46 sites), only energy quality (0.26 [0.03, 0.49]) emerged as a clear predictor of biodiversity, while energy quantity (0.11 [-0.33, 0.51]) and salinity (0.08 [-0.11, 0.28]) had no clear effect.

The importance of energy quality as a predictor of biodiversity further manifests when using taxonomic assignments and restricting the analyses to predominantly pelagic or planktonic consumers (Arthropoda, Chaetognatha, Cnidaria, Chordata, Ctenophora, Rotifera, Amoebozoa, Discosea, Tubulinea, and Bigyra [see Methods]). After applying the incidence-frequency based Chao2 estimator to this taxonomically-informed dataset (see Methods), the global model identified energy quantity (-0.32 [-0.42, −0.22]; 100% probability) and quality (0.16 [0.02, 0.30]; 98.9% probability) as opposing drivers of biodiversity (Fig. 3): while diversity across locations declined with increasing energy quantity, the number of taxa rose as energy quality increased. As previously, the energy quantity effect was mirrored by an opposing, equally strong effect of salinity (0.35 [0.28, 0.42], 100% probability), highlighting the strong, freshwater-mediated correlation between salinity and energy quantity. Finally, these conclusions were supported by effects of energy quantity (-0.37 [-0.51, −0.20]; 99.8% probability) and quality (0.13 [0.01, 0.25]; 97.2% probability) in the regional analysis (Fig. 3). Thus, our results highlight a strong, consistent role of water biogeochemistry on open-water consumer biodiversity: freshwater inflow and its impacts on salinity and associated loads of particulate organic matter modulate coastal species richness, but energy quality independently creates hotspots of planktonic consumer biodiversity in coastal waters at regional and global scales.

**Figure 3.**
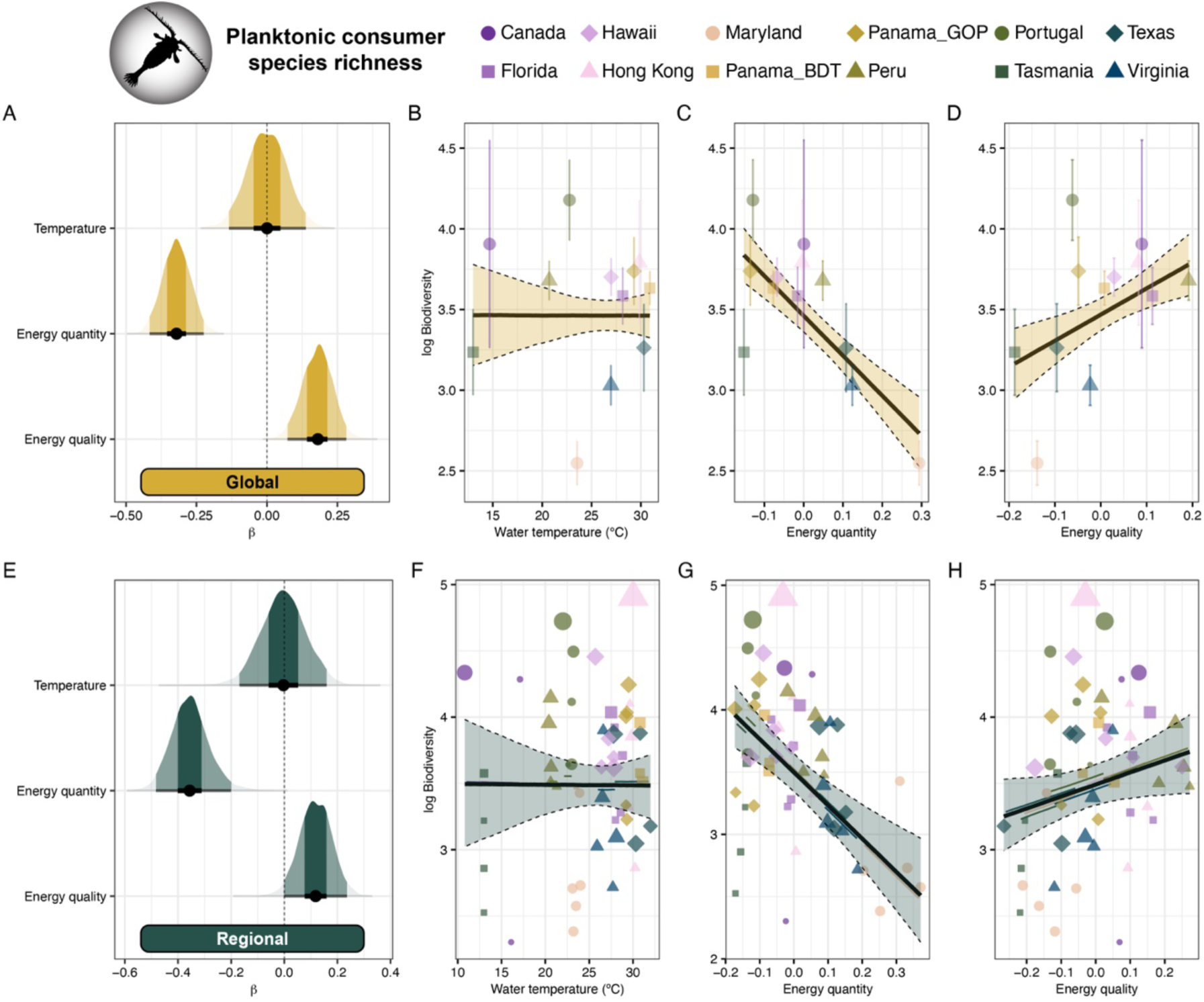
Relationships between the diversity of pelagic or planktonic phyla and environmental predictors across scales. (A and E) Posterior parameter distributions from Bayesian models indicating the strength and direction of scaled predictor effects in the global (A) and regional (E) models. All predictor variables were scaled and centered prior to the analysis for comparability. Shading and black caterpillar bars represent the 50% and 95% credible intervals. (B – D and F – H) Partial effects plots of predictor variables. For the global model (B – D), points and error bars represent the weighted means and standard errors of the Chao2 estimator (based on the lowest identified taxon), while the thick black line and dashed ribbon represent the predicted partial posterior fit derived from the model. For the regional model (F – H), points represent the weighted Chao2 estimator at each site as derived from the replicate samples, with the size of the point reflecting the weight of each observation. The thick black line and dashed ribbon represent the predicted marginalized partial posterior fit derived from the model, while colored, solid lines represent the conditional slopes for each location. Combinations of shapes and colors correspond to the twelve locations (see legend). Panama_GOP = Gulf of Panamá, Panamá, Panama_BDT = Bocas del Toro, Panamá.

The final line of evidence for a species-energy quality relationship in nearshore, open water systems comes from the community composition of heterotrophic organisms. A canonical analysis of principal coordinates (CAP) of consumer orders showed that sites across the twelve locations broadly overlapped with each other, as expected given the fairly coarse taxonomic resolution. However, the sites tended to arrange along a gradient from colder, low salinity sites with high energy quantity to sites with warmer water, high salinity, and high energy quality. Importantly, there was a convergence of some of the most important planktonic consumer taxa, namely calanoid and cyclopoid copepods, hydrozoan jellyfish, decapods, and gobiiform fishes toward sites with the highest energy quality (Fig. 4). Therefore, the availability of fresh, marine-derived, high-quality energy attracts a wide range of important open-water consumer species at local scales, while sites with large quantities of poor-quality energy are depauperate in these taxa and, instead, dominated by more benthic-dwelling taxa (e.g., spionid worms).

**Figure 4.**
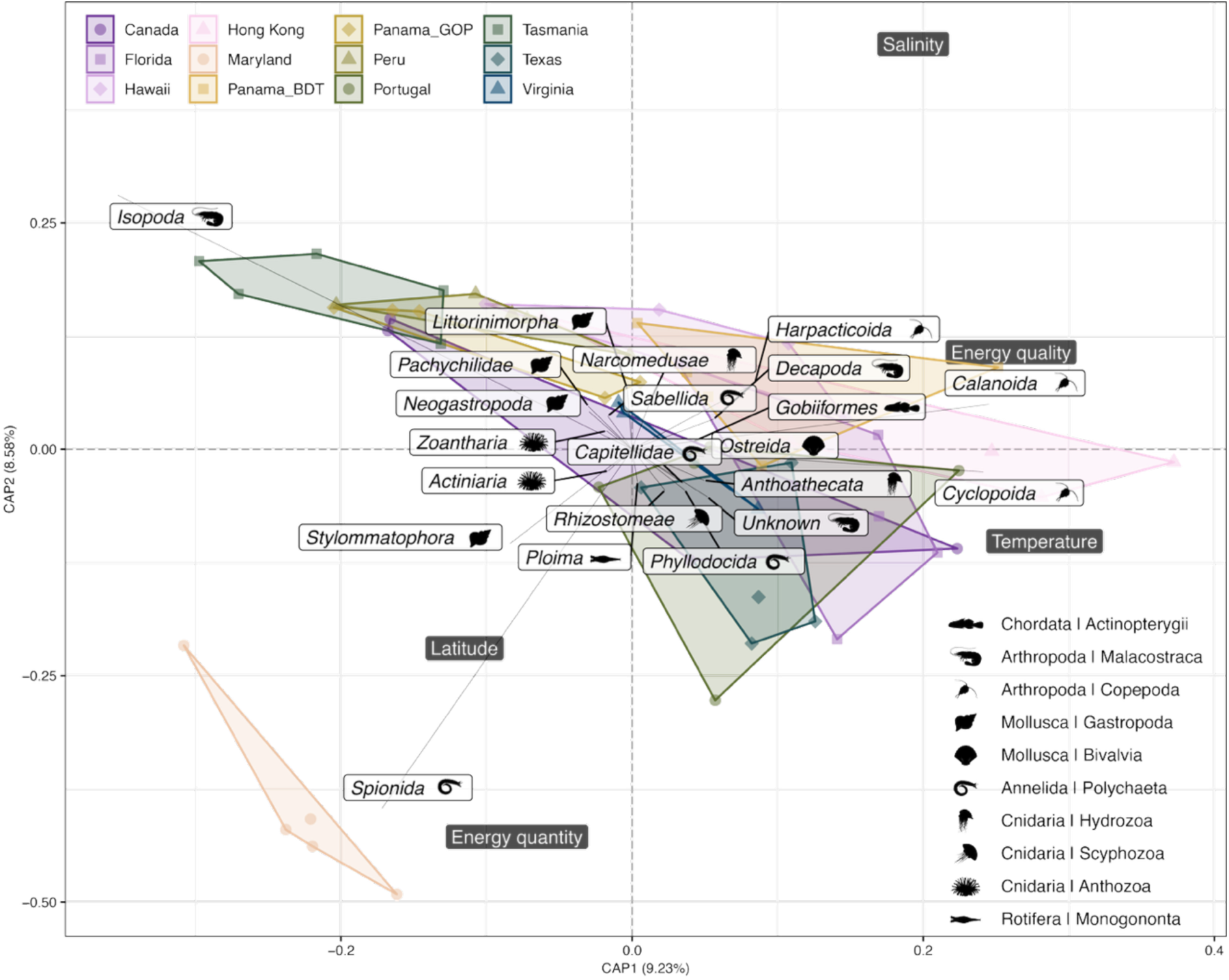
Canonical analysis of principal coordinates (CAP) of the community composition of consumers across the twelve locations. Colored points of different shapes and polygons correspond to the sites and locations, while dark gray labels indicate the positioning of the constraining environmental and biogeochemical variables. White labels with silhouettes represent the taxa driving community separation, performed at the taxonomic level of order and highlighting the ten most extreme taxa on each axis (minimum and maximum). Silhouettes indicate the higher taxonomy for each order. Note that both constraints (divided by 2) and loadings (divided by 3) have been rescaled to facilitate interpretation. Panama_GOP = Gulf of Panamá, Panamá, Panama_BDT = Bocas del Toro, Panamá.

## Discussion

The distribution of life on Earth has long been linked to available energy. We integrate across three ecological paradigms –the species-energy relationship (1), ecological stoichiometry (39), and resource patchiness in the open water (26)– to explain how hotspots of biological activity may arise in nearshore pelagic ecosystems. We first reveal that the environmental filter imposed by low salinity overrides any positive effects of energy availability in carbon-rich waters that are predicted by species-energy theory. While vast amounts of chemical energy enter coastal oceans through freshwater streams, rivers, and estuaries, few organisms can persist in these low salinity environments to harness it. Instead, more saline sites with lower carbon and nitrogen concentrations but fresh, nutrient-rich, marine-derived energy create hotspots of consumer diversity where species, such as important planktonic consumer taxa, congregate. Thus, freshwater inflow and the availability of high-quality energy shape gradients of global and regional diversity in coastal systems and may contribute to the concentration of planktonic consumer species that form the foundation for aggregations of large predators like finfishes, seabirds, and marine mammals. These dynamics may be susceptible to human-mediated changes in coastal ocean biogeochemistry: as intensifying land-use practices and escalating climate change generate intensifying pulses of freshwater laden with recalcitrant terrestrial organic matter, nearshore pools of high-quality energy and their associated diverse consumer communities may become diluted or displaced farther from shore.

### Species–energy relationships across coastal seas

Most of our knowledge about species-energy relationships comes from terrestrial ecosystems, where temperature and other proxies of electromagnetic or chemical energy have generally been successful predictors of species richness.

For example, the quantity of organic material available for consumption (often proxied by vegetation cover, plant biomass, actual evapotranspiration, or the Normalized Difference Vegetation Index) positively correlates with consumer biodiversity, including birds (44), ants (24), beetles (45), mammals (46), and across taxa (47, 48). At face value, our results appear to contradict this paradigm, as energy quantity (in terms of total chemical energy, defined as a latent variable) showed a negative relationship with marine biodiversity. However, this disparity likely stems from three intertwined dynamics that highlight fundamental differences between terrestrial and marine systems and the effect of freshwater runoff in nearshore ecosystems. First, in terrestrial systems, measures of plant productivity, biomass, or vegetation cover are inextricably linked to the three-dimensional complexity these producers provide (49), which bolsters consumer richness (50, 51). Conversely, in open water, primary producers do not offer structural habitat, negating any non-trophic positive effects on consumer richness. Second, since many open water environments are depauperate in nutrients and suspended organic matter, areas with high energy availability are (at the global scale) often located in estuaries or large deltas, where rivers shed large amounts of carbon, nitrogen, phosphorus, and trace elements into the ocean (52). Yet, these fluxes are also tied to fluctuations in water chemistry, chiefly salinity, which creates harsh conditions that few species can withstand, resulting in an environmental filter that creates depauperate species assemblages in transitional estuarine waters (53). Accordingly, salinity emerged as a major driver of biodiversity in our analyses. Finally, the organic matter available in coastal waters that are influenced by terrestrial runoff is often derived from aged, highly recalcitrant resources (e.g., plant-based cellulose or lignin) that are only available for consumption to a few specialized species (54). Therefore, although non-causal in nature (i.e., energy quantity does not directly cause low species richness), high energy availability –when quantified as bulk chemical energy comparable to proxies in terrestrial ecosystems, such as plant biomass or NDVI– correlates with relatively species-poor assemblages in coastal oceans at the global scale. This corroborates evidence from pelagic systems, where, zones of high POM availability due to river outflow often support fewer consumer species than nearby waters with lower POM (34).

Unlike energy quantity, the quality of available energy has rarely been examined as a driver of species-energy relationships in terrestrial or aquatic systems (but see (45, 55, 56)). Yet, it emerged as a robust correlate of biodiversity in our coastal dataset, suggesting that low carbon to nitrogen (C:N) ratios, marine δ^13^C signatures, and high lability of POM are associated with higher consumer diversity. Despite the tight constraints on ocean biogeochemistry (57), POM stoichiometry can vary significantly across locations and seascapes due to the interactive effects of nutrient regimes, producer taxon, turbidity, temperature, and ecological succession in phytoplankton community composition (58–60). Similarly, δ^13^C values in coastal POM are primarily influenced by their source: marine phytoplankton-derived carbon typically hovers around -20‰, while terrestrially-derived carbon is generally more depleted (e.g., < -25‰) (61).

Finally, the lability of POM frequently covaries with these two variables, as the most recalcitrant organic matter is typically derived from terrestrial organic matter with high C:N ratios.

Collectively, this creates a clear energy quality axis that promotes coastal biodiversity, likely due to causal, nutritional pathways. Protein-dense (N-rich), labile resources, such as marine diatoms, generally provide zooplankton with superior nutrition (62), including critical ω-3 polyunsaturated fatty-acids (PUFAs) and sterols without indigestible compounds like lignin or cellulose (63, 64). Therefore, most zooplankton will preferentially use labile, marine-derived phytoplankton across coastal ecotones even with significant influx of terrestrial organic matter (65). While these organism-level preferences are well documented, our results demonstrate that energy quality also predicts local consumer biodiversity independent from well-established diversity gradients across the freshwater-to-marine continuum (53).

### Energy-mediated coastal marine biodiversity hotspots

Global marine biodiversity patterns are governed by a wide range of factors, including biogeography, environmental conditions, and taxon-specific life-histories (66, 67). While ocean biogeochemistry has frequently been used to guide the delineation of biogeographic provinces in the pelagic zone (68, 69), the structure of coastal fauna is typically driven by geological history (70, 71), area (72, 73), and ambient temperature (3, 74). Our results highlight the importance of biogeochemical energy quality for coastal marine biodiversity, especially for taxa with predominantly pelagic or planktonic lifestyles. Energy quality has been considered in the interpretation of global marine productivity patterns but is often overshadowed by resource quantity and has not been linked to consumer biodiversity. For example, the surfacing of nutrient-rich waters in Eastern Boundary Upwelling Systems (EBUS) greatly boosts phytoplankton growth, thus increasing resource availability for pelagic consumers (75). Perhaps more importantly, nitrate (NO3-) upwelling may especially promote the growth of protein-rich phytoplankton species (e.g., diatoms) replete with ω-3 PUFAs and other vital N-rich compounds (76). These nutritionally beneficial phytoplankton species disproportionally increase the biomass and productivity of consumers (77, 78). Thus, the sustained availability of high-quality resources may favor population growth and persistence through physiological benefits, perhaps increasing regional and local species richness over sustained timescales.

Similarly, at the regional scale, variability in species richness is frequently linked to resource availability and productivity. However, especially in aquatic ecosystems, these relationships are often unimodal, where areas of intermediate productivity support the most species (47, 79, 80). While we did not explicitly test for non-linear relationships (see Methods), our results suggest a negative relationship between energy quantity and species richness that is largely mediated by freshwater inflow and its effect on salinity. In contrast, energy quality boosted biodiversity independent of salinity, including for primarily pelagic or planktonic consumers. This result builds on hitherto limited evidence for a role of energy quality in mediating consumer diversity in terrestrial and freshwater consumers. Leaf litter insect assemblages in Central and South America and Southeast Asia exhibit increased biodiversity when resource quality is higher (56, 81, 82), and xylophagous beetles feeding on dead wood showed higher diversity with increasing substratum quality (45). In contrast, for terrestrial plants and stream invertebrates, dominance by otherwise resource limited species reduced assemblage biodiversity in high quality resource environments (55, 83).

In coastal, open-water systems, high quality energy favors higher consumer species richness, especially attracting dominant planktonic consumers such as cyclopoid and calanoid copepods, hydrozoan jellyfish, and gobiiform fishes, which are highly abundant as larvae in nearshore plankton communities and include fully planktonic species (the gobiiform genus *Schindleria*) (84). Similar to pulse-driven leaf-litter communities in tropical rainforests (where decomposition is rapid, nutrient limited, and determined by local resource quality (85, 86)), and deadwood resources (which occur sparsely and unpredictably across landscapes (87)), the patchy and ephemeral availability of high-quality energy in coastal waters prevents the establishment of dominant taxa, thus resulting in transient hotspots of zooplankton biodiversity. While the structuring forces of zooplankton communities may be variable (88–90), greater species richness likely correlates with high zooplankton foraging activity through either niche-based or abundance-driven mechanisms, resulting in increased production of the infochemical dimethyl sulfide (DMS) (91) and the attraction of larger vertebrate predators (92–94). Thus, in nearshore ecosystems, patches of high energy quality, but not quantity, may increase the biodiversity of planktonic taxa and lay the foundation for the emergence of foraging hotspots that support a vast range of vertebrate taxa. The positive relationship between salinity and biodiversity in our coastal study suggests that these hotspots are likely to occur in more saline, oceanic waters; however, the independent effects of energy quality highlight strong spatial variability, which is characteristic of open-water food webs (26).

### Conclusions

Our results suggest a central role of ocean biogeochemistry in coastal marine species-energy relationships, which may influence the patchiness of food resources in open water. Yet, there are three caveats that limit the strengths of our inferences. First, our dataset was limited to twelve coastal locations for which there was strong collinearity between energy quantity and salinity. While this may be largely rooted in a natural, causal relationship (i.e., less saline waters generally exhibit higher energy availability), it is impossible to fully disentangle the roles of energy quantity and salinity across our dataset without broader spatial and temporal coverage, including locations or seasonal conditions with high salinity and energy quantities. Notably, our results were robust to restricting the dataset to locations or sites with strictly marine conditions (salinity >30), suggesting that the observed patterns are not simply due to a few estuarine sites.

Second, our analyses were limited to a single universal primer set (COI). While this was an appropriate marker for our specific question, primer-specific coverage and reference-library completeness vary among taxa and regions, which may bias richness estimates and our obtained relationships (95). Future examinations using complementary markers (e.g., 12S, 16S, 18S, 23S, ITS, etc.) will help resolve the relationships between chemical energy and biodiversity more comprehensively, while also allowing a deeper examination of primary producers, primary consumers, and secondary consumers. Finally, due to the limited number of samples within locations and at global scales (N = 5 and N = 12, respectively), we were unable to robustly test for non-linear fits in species-energy relationships.

Nevertheless, if salinity and energy quality jointly govern the occurrence of hotspots of high zooplankton biodiversity in coastal oceans, then the increasingly erratic and intense patterns of terrestrial run-off into nearshore ecosystems may threaten the ecological dynamics that provide irreplaceable services to humanity. Continental run-off has increased in the past century due to climate change and intensifying land-use practices (96). While freshwater inflow is a crucial determinant of estuarine ecosystem functioning, excessive pulses of carbon-laden, poorly managed runoff can negatively affect communities in nearshore ecosystems through physiological stress and the dominance of recalcitrant, land-derived organic matter that is inaccessible to most consumers (97–99). Our results suggest that the combined impacts of freshwater and recalcitrant, carbon-rich, land-derived organic matter will independently reduce zooplankton biodiversity in coastal seascapes. As a consequence, hotspots of nearshore biodiversity are likely to become sparser, which may compromise two benefits that coastal oceans provide to humanity: the supply of blue foods and the support of marine recreational activities such as whale watching, birding, and recreational fishing (100). While the quality of energetic resources is difficult to observe and quantify, it plays a critical but vulnerable role in governing planktonic consumer biodiversity and foraging dynamics in nearshore open water ecosystems.

## Materials and Methods

### Sample collection

We implemented biogeochemistry and eDNA surveys across twelve coastal locations from May 30^th^ to July 19^th^ 2023. Representatives from partner institutions associated with each location received identical kits for water filtration following a standardized protocol that included proper sterilization of utensils and workspaces. Within each location, we sampled five sites that were separated by at least 1km at depths between 1m and 10m. Within each site, three replicate water samples were taken for biogeochemistry (*N* = 180), and five replicate water samples were obtained with sterilized equipment for eDNA analysis (*N* = 300). At each site, we included one 1L blank sterile water sample (Milli-Q or distilled water; *N* = 60). In addition, salinity and temperature were measured at each site using handheld sondes. For Tasmania, no site-specific temperature and salinity data were collected, which were instead imputed from the HYCOM model and cross-checked by local experts.

### Biogeochemistry

Water samples for biogeochemistry were filtered through 47 mm 0.7 µm glass fiber filters, which were combusted to remove carbon prior to filtering. The volume of filtered water ranged from 100 mL to 1L, depending on the time it took for filters to become clogged. Prior to shipping, biogeochemistry filters were dried at 60°C for 24 hours then placed in aluminum foil. All dried filters were shipped to the University of Texas Austin Marine Science Institute (UTMSI) and stored at room temperature until processing.

At UTMSI, filters were either quartered or halved depending on their color (as a proxy for their concentration) using a methanol rinsed scalpel, then fumigated for 24 hours with 12N hydrochloric acid to remove inorganic carbon. Filters were removed from fumigation and left to dry out for approximately 1 hour. Filters were then folded and wrapped in tin capsules (Costech, USA). Samples were analyzed for Particulate Organic Carbon (POC, as μmol C L^-1^ of seawater), Particulate Organic Nitrogen (PON, as μmol N L^-1^ of seawater), δ^15^N, and δ^13^C at UTMSI using Carlo Erba NC 2500 Elemental Analyzer interfaced through a Conflo II to a Finnigan MAT (Thermo) Delta Plus XL isotopic ratio mass spectrometer. Stable isotopes are reported relative to international standards of Vienna Pee Dee Belemnite (C) and air (N). Final values are calculated using the equation: δ [isotope] = [(Rsample/ Rstandard) − 1], where R is the ratio of the stable isotope to its natural form (^13^C/^12^C or ^15^N/^14^N), and units are permille. Precision for the C or N content was within 5% and 0.2‰ for δ^13^C and δ^15^N, respectively.

Total hydrolyzable amino acids (THAAs) were extracted from the filters following Liu and Xue (2020) (41). Two punches (1 cm diameter) were taken from each filter and combined in a glass vial containing 2 mL of 6 N hydrochloric acid. The vial was purged with nitrogen gas, sealed, vortexed, and hydrolyzed for 20 hours at 110°C. Contents were then filtered using glass wool, dried under nitrogen gas and stored at 4°C until analysis. Samples were rehydrated with HPLC-grade deionized water and methanol. Sodium acetate (0.05 M, pH 5.7) and tetrahydrofuran (5%) formed eluent A, with methanol as eluent B. The method ramped from 20% to 50% eluent B in 40 minutes, then to 100% eluent B in 20 minutes. After derivatization by o-phthalaldehyde, 16 amino acids were detected by fluorescence: aspartic acid (ASP), glutamic acid (GLU), histidine (HIS), serine (SER), arginine (ARG), glycine (GLY), threonine (THR), β-alanine (BALA), alanine (ALA), tyrosine (TYR), γ-aminobutyric acid (GABA), methionine (MET), valine (VAL), phenylalanine (PHE), isoleucine (ILE), and leucine (LEU). The identification and concentration of these amino acids were determined via comparison of peak retention times to a certified standard mix (Sigma). Duplicate analyses were within 10% of obtained values, indicating robust analytical precision. The contribution of amino acids to organic carbon (THAA) was calculated for each sample and added as a variable in uMol C/L.

### Environmental DNA

Here, we define environmental DNA (*e*DNA) as all DNA recovered from bulk water samples, including free (extra-organismal), particle-associated, and whole-organism DNA captured on the filter. Water samples for eDNA were filtered through 47 mm 1.5 µm glass fiber filters using a vacuum pump. Water was filtered until the filter clogged, with volumes ranging from 140ml to 3L. The volume of filtered water did not affect biodiversity estimates, as evident from the two locations where different sample volumes were obtained across the five sites (Fig. S2). After filtration, filters were cut in half with one half placed in Powerbead Pro tubes with 600µL CD1 lysis buffer (materials and reagents from Qiagen DNeasy PowerSoil Pro Kit) and the other half frozen at -20 or -80°C for long-term storage. The CD1-preserved filters were shipped to UTMSI and stored at room temperature until processing.

DNA extractions were performed at UTMSI in a fume hood under sterile conditions. We used the DNeasy PowerSoil Pro Kit (Qiagen) following the manufacturer’s protocol, beginning with the homogenization step since the samples were already stored in the lysis buffer (CD1 solution).

Negative controls were included to account for potential contamination during extraction. DNA extracts were quantified using a Qubit Fluorometer (Invitrogen) and diluted to equal concentrations (∼4 ng/ul) prior to PCR amplification. DNA extracts then underwent PCR amplification, targeting the COI gene region (313 bp) using universal metazoan primers: forward 5’-GGWACWGGWTGAACWGTWTAYCCYCC-3’; reverse 5’-TAIACYTCIGGRTGICCRAARAAYCA-3’ (101, 102). Reactions were made to 12 µL using a mixture of 6 µL AmpliTaq Gold 360 Master Mix (ThermoFisher Scientific), 0.7µL of each primer, 2.6µL of nuclease free water, and 2µL DNA extract. Thermocycler PCR conditions were as follows: 94°C for 10 minutes; followed by 16 initial cycles: denaturation for 10 seconds at 94°C, annealing for 30 seconds at 62°C (−1°C per cycle), and extension for 60 seconds at 72°C; followed by 25 cycles: denaturation for 10 seconds at 94°C, annealing at 46°C for 30 seconds, and extension for 30 seconds at 72°C; with a final extension of 72°C for 7 minutes. Amplified products were confirmed using gel electrophoresis with MIDORI Green stain (Nippon Genetics) and run on 1% agarose for 30 minutes at 120V.

PCR products were diluted 1:20 and sent to the Hubbard Centre for Genome Studies (University of New Hampshire) for library preparation and sequencing. Library prep was conducted using a dual-indexing approach with the Nextera XT Index Kit (Illumina). Indexing PCR reactions were carried out in 50 μl volumes containing 5 μl each of Index 1 (i7) and Index 2 (i5), 25 μl of 2× KAPA HiFi HotStart ReadyMix, 10 μl PCR-grade water, and 5 μl DNA template. Thermocycling conditions matched the initial amplification: 95 °C for 3 min; 8 cycles of 95 °C for 30 s, 55 °C for 30 s, and 72 °C for 30 s; and a final extension at 72 °C for 5 min. Final libraries were cleaned with AMPure XP Beads and quantified using a Qubit Fluorometer (Invitrogen). All samples were pooled, normalized to 4 nM, and size selected using a Pippin Prep before sequencing on an Illumina NovaSeq 6000 using a 500-cycle NovaSeq Reagent Kit (Illumina).

### Bioinformatics and taxonomic assignments

Raw paired-end reads were processed with QIIME 2 (2024.5) (103). Primers were removed with the QIIME2 cutadapt trim-paired plugin (104) and reads were denoised using the dada2 denoise-paired plugin (105). Truncation lengths for forward and reverse reads were chosen based on interactive quality plots (qiime demux summarize) to remove low-quality tails. Reads were then filtered based on the default expected error threshold (maxEE = 2), and the default DADA2 consensus method for chimera removal. Following independent denoising, we exported ASV sequences.

We assigned taxonomy using BLAST+ (v2.16.0) (106) against a custom merged reference database including MIDORI2 (GB265; (107)), the Barcode of Life Data System (BOLD; (108)), and two personal, curated databases of marine invertebrates. The merged reference database was standardized to BOLD taxonomic conventions, de-duplicated, and restricted to marine taxa by curating a list of exclusively terrestrial taxa and filtering the database against this list (see Table S1). This was done due to the overwhelming prevalence of terrestrial arthropods in the reference library, resulting in a large quantity of unrealistic assignments to terrestrial taxa. BLAST outputs were filtered based on taxon-specific standardized confidence thresholds using the Error-Based BLAST Sequence (EBBS) R shiny application (109) with the following settings: minimum alignment length 250 bp, maximum e-value 0.01, minimum query coverage 80%, “best-shared” assignment (LCA-like), and taxon-specific confidence thresholds targeting a 10% error rate. Downstream curation in R removed ASVs represented by less than 3 total reads (singletons and doubletons), sequences detected in negative controls, and samples with less than 1,000 reads to mitigate variations in sequence depths. Using the taxonomic assignments, we summed the number of reads for each taxon at the lowest taxonomic level for each sample, resulting in a total of 1,141 taxa. More than 60% of taxonomic identifications were made to at least the genus level using taxon-specific thresholds following (109).

To curate a dataset with only primarily planktonic or pelagic consumer taxa, we filtered the dataset at the phylum level, retaining only ASVs assigned to the following phyla: Arthropoda, Amoebozoa, Bigyra, Chordata, Chaetognatha, Cnidaria, Ctenophora, Discosea, Rotifera, and Tubulinea. While some of these phyla (e.g., Arthropoda) include a large number of non-planktonic organisms, quantifications of taxonomic representation revealed that 40.2% of sequences were assigned to Copepoda, while 47.3% were assigned to Malacostraca, with amphipods (31.2%) and decapods (40.0%) as the dominant orders within Malacostraca. This suggests that, while arthropods are a highly diverse phylum, our data mostly contained taxa that are planktonic consumers (although decapods and amphipods also include many benthic taxa). We included single-celled plankton-like ciliates and amoebas due to their important role as primary consumers in planktonic communities, but their representation in the dataset was limited (<10% of taxa in the final dataset). The total planktonic consumer dataset contained 322 taxa, of which almost 90% (89.1%) consisted of arthropods, chordates, cnidarians, and chaetognaths (in declining order of their representation). Finally, we removed samples that only had a single species present based on the reduced dataset.

### Statistical analyses

For the biogeochemistry dataset, we first used the amino acid data to calculate a Degradation Index (DI) via a Principal Component Analysis (PCA) of the molar percent contribution of amino acids to the amount of total hydrolyzable amino acids (THAA). Each amino acid was converted to mole percent (molbiomarker/moltotal), and the PCA was run on the scaled and centered concentrations of each amino acid. We then plotted the scores and vectors over the first two PC-axes, which explained 21.1% and 17.2% of the variability in the water samples, respectively (Fig. S3). Following Xue et al. (110) and Liu and Xue (41), the orientation of amino acid degradation products (primarily β-alanine [BALA] and γ-aminobutyric acid [GABA]) vs. amino acids indicative of fresh organic matter (primarily aspartic acid [ASP] and glutamic acid [GLU]) were used to interpret the direction of the degradation index (DI) axis to our data. PCA scores along this axis were then extracted for each sample to construct the DI, in our case identifying samples with low scores as more degraded, while samples with higher scores on PC1 can be interpreted as containing fresher organic matter. Since there was also significant spread along the second PC-axis, we also extracted the scores on PC2 despite the lack of a clearly defined biogeochemical differentiation along that axis.

We then integrated the degradation index value into the broader biogeochemistry dataset to obtain latent variables indicative of overall seawater biogeochemistry, as well as energy quantity and quality across samples. This was done due to strong collinearity among several of the biogeochemical variables (e.g., μmol C L^-1^, μmol N L^-1^, and %THAA) and significant collinearity between these same variables and salinity (in PSU, practical salinity units). Specifically, we conducted a second PCA that included the following variables: μmol C L^-1^, μmol N L^-1^, %THAA, C:N ratio, degradation index (PC1), degradation index (PC2), δ^15^N, and δ^13^C. We again scaled and centered each variable and plotted the resulting ordination across the two principal components, which explained 43.9% and 18.9% of the variability across PC1 and PC2, respectively. To obtain values for this latent ‘energy index’, we again extracted scores for each sample on PC1 and PC2. Based on their loadings (Fig. 1B), these axes were henceforth denoted as “energy quantity” (PC1) and “energy quality” (PC2), and both were multiplied by -1 to transform their values to be intuitive (low values indicating low energy quantity and quality, respectively). Note that these indices are naturally imperfect: they do not measure caloric potential or nutritional value in a true biochemical or nutritional sense, but instead represent proxies of the broader resource environment that species have at their disposal. As such, our approach is comparable to a vast number of studies that use proxies of energy availability (e.g., temperature, latitude, tree cover, or NDVI). Using univariate variables (e.g., μmol C L^-1^, μmol N L^-1^, C:N ratio, or the degradation index) instead of the derived latent variables in subsequent analyses did not change any of our conclusions but introduced substantial limitations due to collinearity (see Supplementary Materials).

To obtain diversity estimates, we used the incidence-frequency based Chao2 estimator (42) across the five eDNA water samples taken at each site, resulting in coverage-based asymptotic estimates of ASV-richness (taxonomically-uninformed dataset) and species richness of planktonic consumers (taxonomically-informed dataset) along with their uncertainty estimates for each site (42). This dual approach (with and without taxonomy) was performed since many ASVs in our dataset were not reliably assigned to marine taxa, which likely reflects the incompleteness of COI reference libraries for many marine taxa (111). We used both the full scope of the metabarcoding dataset (irrespective of the unknown identities of the ASVs) and a more refined, targeted dataset of assigned taxonomic identities to species putatively most affected by coastal marine POM (heterotrophic taxa with predominantly pelagic or planktonic life histories). We then applied a meta-analysis workflow to the biodiversity data to account for uncertainty in the Chao2 biodiversity estimates, computing log-transformed richness estimates and inverse-variance weights for each site (112). Specifically, for each site *i*, the log-transformed richness estimate is:

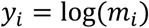

where *m_i_* is the Chao2 species richness estimate. The approximate variance of *y_i_* on the log-scale was then obtained using the delta method applied to the arithmetic-scale standard error *si* of the Chao2 estimator:

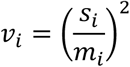

Where *v_i_* is the variance of *y_i_* on the log-scale and *s_i_* is the standard error of the Chao2 estimator on the arithmetic scale. The weight for each observation is then the reciprocal of this log-scale variance:

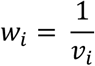

Weights were then normalized across all observations so that their mean equals 1:

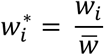

Where 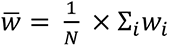, i.e., the mean of the unnormalized weights across *N* observations. This mean-normalization preserved the relative weighting structure, while ensuring their mean is centered around 1, a step that was necessary for stable likelihood calculation in the Bayesian models at the regional scale. We then merged the biogeochemistry dataset with the two eDNA datasets (ASVs and taxonomically-informed planktonic consumers), retaining average values for biogeochemical indicators and estimated richness values for each site. For the reduced dataset of planktonic consumers, we removed one site (Texas Site 1) since its low richness and uncertainty rendered it incompatible with the weighted regression analyses.

Using the combined datasets, we ran six different analyses to examine drivers of biodiversity in coastal oceans across locations (global model) and within locations (regional model), each applied to the ASV-richness estimates and the taxonomically-informed planktonic consumer dataset. Each global model was run twice using either energy quantity, quality, and temperature or only salinity as predictors due to the strong collinearity between salinity and energy quantity. Given significant spatial autocorrelation among sites within locations (Moran test: Moran I statistic = 0.47, *p* = 0.001), we summarized information at the location level for all global models. Specifically, using the ASV or planktonic consumer richness estimates for each location based on the five sites and their weights, we calculated the weighted means, weighted variances, effective sample sizes, and finally, weighted standard errors. For observations *y_i_* with weights *w_i_* (as derived above), the mean for each location *j* is:

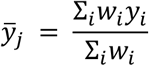

The weighted variance of log-richness is:

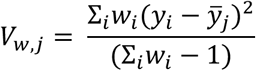

The effective sample size is:

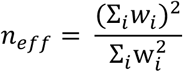

And the weighted standard error of mean log-richness as:

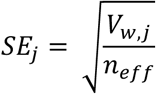

Using these values, we ran two Bayesian linear models with the weighted mean richness estimate as the response and the weighted standard error specified as measurement error on this term, essentially treating the mean richness estimate as the measured variable (*R_obs_*) with a known measurement error (*R_err_*), defined as *R_obs_*_,*i*_ ∼ N(*R_true_*_,*i*_, *R_err_*_,*i*_). This allowed us to retain the uncertainty around the response variable and its weighted structure, while modeling its relationship with our chosen predictors. Given the normal distribution of log-transformed richness values, we used a Gaussian error distribution. In the first model, we fitted the z-transformed (i.e., scaled and centered) mean values of water temperature (in °C), energy quantity, and energy quality for each location as predictors (biogeochemistry model), while also fitting a model with only salinity (salinity model). We performed the same procedure for both the ASV-richness dataset and the taxonomically-informed planktonic consumer dataset. We ran these models using *brms* default uninformative priors on the intercept (113), while specifying weakly-informative priors on the slope parameters (student-t with 3 degrees of freedom, location = 0, and scale = 1) to aid model convergence. This resulted in the following model structure:

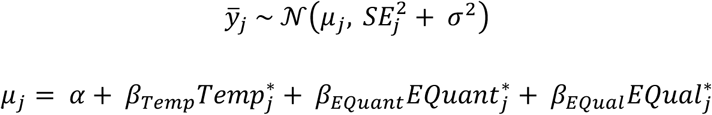

Where *ȳ_j_* is the weighted mean of log-richness for each location *j*, 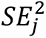 is the squared weighted standard error of that mean (i.e., 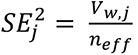, the estimated measurement variance of *ȳ_j_* on the log scale), *σ*^2^ is the residual process variance, and 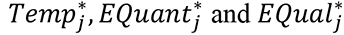 are the scaled values of water temperature, energy quantity, and energy quality, respectively, with the priors for the slope estimates specified as:

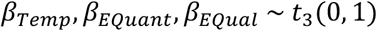

The same structure was retained for the salinity model, but with only salinity (*Sal*) as the predictor.

For the regional models, we ran Bayesian linear mixed models in which we used the site-level estimates of richness as the response variable and specified the calculated mean-normalized weights (as specified above) as a multiplier on the likelihood contributions of each observation.

Mean-normalization was necessary for these models because *brms* implements weights as a direct precision multiplier on the likelihood. Therefore, weights with very small values would dramatically inflate the residual variance and allow the prior to dominate the posterior. We then fitted temperature, energy quantity, and energy quality as fixed effects, while specifying location as a random intercept. We again used a Gaussian error distribution due to the near-normal distribution of log-transformed richness estimates. We omitted salinity from this model due to the high collinearity with energy quantity and did not run dedicated models for salinity in this case.

This approach permitted us to derive the average shape of the relationship between richness and the different covariates across the different locations based on the local spread of values. While it assumes a fixed slope for this relationship among the locations, we were unable to run a reliable random-slope model (as ascertained by tests of chain convergence and posterior predictive checks). The same model was run for both the ASV-richness dataset and the taxonomically-informed planktonic consumer dataset. We again used default non-informative priors on the intercept and the variance estimates, and weakly informative student-t priors on the slope estimates. For this model type, the structure for each site-level observation *k* in location *l*[*k*] was:

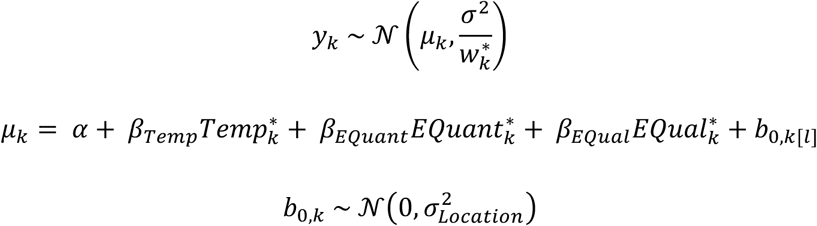

Where 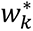 is the mean-normalized inverse-variance (so that a site with a more precise richness estimate receives a smaller effective residual variance, and thus contributes more to the likelihood), and *b*_<,*k*_ is the random intercept for each location. Finally, due to the potential confounding mechanisms arising from the backdoor criterion (114), we also re-ran the two regional analyses with Location added as a fixed effect. The outcomes from this analysis were quantitatively and qualitatively indistinguishable from the other two models. All models in this study were run for 20,000 iterations with a warmup of 10,000 iterations and a thinning rate of 10. All models were run with four chains, and we assessed model performance using posterior predictive checks, Rhat values, and visual inspections of chain convergence (115).

Finally, we used the taxonomically-informed dataset across consumer groups (i.e., not restricted to planktonic taxa but without taxa generally classified as primary consumers) to assess community composition of consumers across locations and sites and the dominant environmental constraints. Specifically, we first removed all reads that were assigned to primary producers (e.g., phytoplankton, macroalgae) or fungi, resulting in a dataset of 668 operational taxonomic units.

We then summarized taxa at the order level and performed a Canonical Analysis of Principal Coordinates (CAP) using the Bray-Curtis dissimilarity on the Hellinger-transformed sequence reads in each taxon across sites. We fitted absolute latitude, temperature, salinity, energy quantity, and energy quality as constraints on the ordination and then plotted the ordination. To examine the taxa that characterized community composition across the environmental constraints, we selected the ten taxa with the highest and lowest scores on the two orthogonal axes and plotted them in the same ordination space. All statistical analyses were performed in R (116), using the packages *brms* (113), *metafor* (112), *vegan* (117), and *iNEXT* (42) for analyses, as well as the *tidyverse* (118) package and various ancillary packages for data processing and visualization.

## Acknowledgments

We thank all volunteers who helped with water collections across the network, including Jesse J. Carter, Paepae o Heʻeia, Kaleonani Hurley, Noʻeau Machado, Kyle Bosworth, and Lani Musselman. This work was supported by the Smithsonian Institution’s MarineGEO BEACON Network Project (Biodiversity and Energy Availability across a Coastal Ocean Network). SJB was supported by a National Academies of Science, Engineering and Medicine Early-Career Gulf Research Fellowship. This is contribution number X from the Smithsonian’s MarineGEO and Tennenbaum Marine Observatories Network.

## Classification

Biological Sciences | Ecology

## References

1. D. H. Wright, Species-energy theory: an extension of species-area theory. Oikos 41, 496–506 (1983).

2. K. J. Gaston, Global patterns in biodiversity. Nature 405, 220–227 (2000).

3. H. Hillebrand, On the generality of the latitudinal diversity gradient. The American Naturalist 163, 192–211 (2004).

4. G. E. Hutchinson, Homage to Santa Rosalia or why are there so many kinds of animals? The American Naturalist 93, 145–159 (1959).

5. A. H. Hurlbert, Species–energy relationships and habitat complexity in bird communities. Ecology Letters 7, 714–720 (2004).

6. D. J. Currie, Energy and large-scale patterns of animal-and plant-species richness. The American Naturalist 137, 27–49 (1991).

7. M. Kaspari, P. S. Ward, M. Yuan, Energy gradients and the geographic distribution of local ant diversity. Oecologia 140, 407–413 (2004).

8. J. T. Kerr, R. Vincent, D. J. Currie, Lepidopteran richness patterns in North America. Ecoscience 5, 448–453 (1998).

9. K. Roy, D. Jablonski, J. W. Valentine, G. Rosenberg, Marine latitudinal diversity gradients: tests of causal hypotheses. Proceedings of the national Academy of Sciences 95, 3699–3702 (1998).

10. S. Rutherford, S. D’Hondt, W. Prell, Environmental controls on the geographic distribution of zooplankton diversity. Nature 400, 749–753 (1999).

11. G. G. Mittelbach, et al., Evolution and the latitudinal diversity gradient: speciation, extinction and biogeography. Ecology letters 10, 315–331 (2007).

12. J. H. Brown, Why are there so many species in the tropics? Journal of biogeography 41, 8–22 (2014).

13. M. T. P. Coelho, et al., Consistent energy-diversity relationships in terrestrial vertebrates. Science 389, 53–57 (2025).

14. J. W. Valentine, D. Jablonski, A twofold role for global energy gradients in marine biodiversity trends. Journal of Biogeography 42, 997–1005 (2015).

15. J. E. Duffy, et al., A Pleistocene legacy structures variation in modern seagrass ecosystems. Proceedings of the National Academy of Sciences 119, e2121425119 (2022).

16. X. Irigoien, J. Huisman, R. P. Harris, Global biodiversity patterns of marine phytoplankton and zooplankton. Nature 429, 863–867 (2004).

17. P. G. Falkowski, R. T. Barber, V. Smetacek, Biogeochemical controls and feedbacks on ocean primary production. science 281, 200–206 (1998).

18. P. Cermeño, F. G. Figueiras, Resource levels, allometric scaling of population abundance, and marine phytoplankton diversity. Limnology and Oceanography 53, 312–321 (2008).

19. A. D. Barton, S. Dutkiewicz, G. Flierl, J. Bragg, M. J. Follows, Patterns of diversity in marine phytoplankton. Science 327, 1509–1511 (2010).

20. D. Righetti, M. Vogt, N. Gruber, A. Psomas, N. E. Zimmermann, Global pattern of phytoplankton diversity driven by temperature and environmental variability. Science Advances 5, eaau6253 (2019).

21. W. F. Ruddiman, Recent planktonic foraminifera: dominance and diversity in North Atlantic surface sediments. Science 164, 1164–1167 (1969).

22. I. Rombouts, et al., Global latitudinal variations in marine copepod diversity and environmental factors. Proceedings of the Royal Society B: Biological Sciences 276, 3053–3062 (2009).

23. R. S. Woodd-Walker, P. Ward, A. Clarke, Large-scale patterns in diversity and community structure of surface water copepods from the Atlantic Ocean. Marine Ecology Progress Series 236, 189–203 (2002).

24. M. Kaspari, S. O’Donnell, J. R. Kercher, Energy, density, and constraints to species richness: ant assemblages along a productivity gradient. The American Naturalist 155, 280–293 (2000).

25. C. Brown, et al., Effects of climate-driven primary production change on marine food webs: implications for fisheries and conservation. Global Change Biology 16, 1194–1212 (2010).

26. K. J. Benoit-Bird, Resource patchiness as a resolution to the food paradox in the sea. The American Naturalist 203, 1–13 (2024).

27. B. Scott, et al., Sub-surface hotspots in shallow seas: fine-scale limited locations of top predator foraging habitat indicated by tidal mixing and sub-surface chlorophyll. Marine Ecology Progress Series 408, 207–226 (2010).

28. K. J. Benoit-Bird, et al., Prey patch patterns predict habitat use by top marine predators with diverse foraging strategies. PloS one 8, e53348 (2013).

29. C. Lambert, et al., Energyscapes pinpoint marine megafauna feeding hotspots in the Mediterranean. Proceedings of the National Academy of Sciences 122, e2412845122 (2025).

30. D. K. Wingfield, et al., The making of a productivity hotspot in the coastal ocean. PloS one 6, e27874 (2011).

31. B. A. Block, et al., Tracking apex marine predator movements in a dynamic ocean. Nature 475, 86– 90 (2011).

32. F. Ribalet, et al., Unveiling a phytoplankton hotspot at a narrow boundary between coastal and offshore waters. Proceedings of the National Academy of Sciences 107, 16571–16576 (2010).

33. S. L. Chown, K. J. Gaston, Patterns in procellariiform diversity as a test of species-energy theory in marine systems. Evolutionary Ecology Research 1, 365–373 (1999).

34. L. Thiers, et al., Combining methods to describe important marine habitats for top predators: Application to identify biological hotspots in tropical waters. PloS one 9, e115057 (2014).

35. B. Guenet, M. Danger, L. Abbadie, G. Lacroix, Priming effect: bridging the gap between terrestrial and aquatic ecology. Ecology 91, 2850–2861 (2010).

36. T. S. Bianchi, The role of terrestrially derived organic carbon in the coastal ocean: A changing paradigm and the priming effect. Proceedings of the National Academy of Sciences 108, 19473– 19481 (2011).

37. J. H. Thorp, M. D. Delong, Dominance of autochthonous autotrophic carbon in food webs of heterotrophic rivers. Oikos 96, 543–550 (2002).

38. A. Pusceddu, A. Dell’Anno, M. Fabiano, R. Danovaro, Quantity and bioavailability of sediment organic matter as signatures of benthic trophic status. Marine Ecology Progress Series 375, 41–52 (2009).

39. R. W. Sterner, J. J. Elser, Ecological stoichiometry: the biology of elements from molecules to the biosphere (Princeton university press, 2003).

40. S. Mäkelin, et al., Linking Resource Quality and Biodiversity to Benthic Ecosystem Functions Across a Land-to-Sea Gradient. Ecosystems 27, 329–345 (2024).

41. Z. Liu, J. Xue, The lability and source of particulate organic matter in the northern gulf of mexico hypoxic zone. Journal of Geophysical Research: Biogeosciences 125, e2020JG005653 (2020).

42. T. Hsieh, K. Ma, A. Chao, iNEXT: an R package for rarefaction and extrapolation of species diversity (H ill numbers). Methods in Ecology and Evolution 7, 1451–1456 (2016).

43. A. Chao, et al., Rarefaction and extrapolation with Hill numbers: a framework for sampling and estimation in species diversity studies. Ecological monographs 84, 45–67 (2014).

44. W. Jetz, C. Rahbek, Geographic range size and determinants of avian species richness. Science 297, 1548–1551 (2002).

45. P. Kriegel, et al., Ambient and substrate energy influence decomposer diversity differentially across trophic levels. Ecology Letters 26, 1157–1173 (2023).

46. W. Jetz, H. Kreft, G. Ceballos, J. Mutke, Global associations between terrestrial producer and vertebrate consumer diversity. Proceedings of the Royal Society B: Biological Sciences 276, 269– 278 (2009).

47. G. G. Mittelbach, et al., What is the observed relationship between species richness and productivity? Ecology 82, 2381–2396 (2001).

48. D. J. Currie, et al., Predictions and tests of climate-based hypotheses of broad-scale variation in taxonomic richness. Ecology letters 7, 1121–1134 (2004).

49. E. A. LaRue, et al., A theoretical framework for the ecological role of three-dimensional structural diversity. Frontiers in Ecology and the Environment 21, 4–13 (2023).

50. R. H. MacArthur, J. W. MacArthur, J. Preer, On bird species diversity. II. Prediction of bird census from habitat measurements. The American Naturalist 96, 167–174 (1962).

51. D. S. Srivastava, Habitat structure, trophic structure and ecosystem function: interactive effects in a bromeliad–insect community. Oecologia 149, 493–504 (2006).

52. R. Chester, “Nutrients, organic carbon and the carbon cycle in sea water” in Marine Geochemistry, (Springer, 1990), pp. 272–320.

53. A. Whitfield, M. Elliott, A. Basset, S. Blaber, R. West, Paradigms in estuarine ecology–a review of the Remane diagram with a suggested revised model for estuaries. Estuarine, Coastal and Shelf Science 97, 78–90 (2012).

54. P. A. Raymond, J. E. Bauer, Riverine export of aged terrestrial organic matter to the North Atlantic Ocean. Nature 409, 497–500 (2001).

55. B. B. Tumolo, S. M. Collins, Y. Guan, A. C. Krist, Resource quantity and quality differentially control stream invertebrate biodiversity across spatial scales. Ecology Letters 26, 2077–2086 (2023).

56. M. Jochum, et al., Resource stoichiometry and availability modulate species richness and biomass of tropical litter macro-invertebrates. Journal of Animal Ecology 86, 1114–1123 (2017).

57. A. C. Redfield, On the proportions of organic derivatives in sea water and their relation to the composition of plankton (University press of Liverpool Liverpool, 1934).

58. T. Tanioka, K. Matsumoto, A meta-analysis on environmental drivers of marine phytoplankton C: N: P. Biogeosciences 17, 2939–2954 (2020).

59. A. Martiny, J. A. Vrugt, F. W. Primeau, M. W. Lomas, Regional variation in the particulate organic carbon to nitrogen ratio in the surface ocean. Global Biogeochemical Cycles 27, 723–731 (2013).

60. A. C. Martiny, et al., Strong latitudinal patterns in the elemental ratios of marine plankton and organic matter. Nature Geoscience 6, 279–283 (2013).

61. M.-T. Verwega, et al., Description of a global marine particulate organic carbon-13 isotope data set. Earth System Science Data 13, 4861–4880 (2021).

62. E. T. Peltomaa, S. L. Aalto, K. M. Vuorio, S. J. Taipale, The importance of phytoplankton biomolecule availability for secondary production. Frontiers in Ecology and Evolution 5, 128 (2017).

63. S. M. Colombo, A. Wacker, C. C. Parrish, M. J. Kainz, M. T. Arts, A fundamental dichotomy in long-chain polyunsaturated fatty acid abundance between and within marine and terrestrial ecosystems. Environmental Reviews 25, 163–174 (2017).

64. A. W. Galloway, M. Winder, Partitioning the relative importance of phylogeny and environmental conditions on phytoplankton fatty acids. PLoS One 10, e0130053 (2015).

65. T. A. Schlacher, R. M. Connolly, A. J. Skillington, T. F. Gaston, Can export of organic matter from estuaries support zooplankton in nearshore, marine plumes? Aquatic Ecology 43, 383–393 (2009).

66. B. W. Bowen, et al., Comparative phylogeography of the ocean planet. Proceedings of the National Academy of Sciences 113, 7962–7969 (2016).

67. D. P. Tittensor, et al., Global patterns and predictors of marine biodiversity across taxa. Nature 466, 1098–1101 (2010).

68. G. Reygondeau, et al., Dynamic biogeochemical provinces in the global ocean. Global Biogeochemical Cycles 27, 1046–1058 (2013).

69. M. D. Spalding, et al., Marine ecoregions of the world: a bioregionalization of coastal and shelf areas. BioScience 57, 573–583 (2007).

70. E. Di Martino, J. B. Jackson, P. D. Taylor, K. G. Johnson, Differences in extinction rates drove modern biogeographic patterns of tropical marine biodiversity. Science Advances 4, eaaq1508 (2018).

71. D. Bellwood, P. Wainwright, C. Fulton, A. Hoey, Assembly rules and functional groups at global biogeographical scales. Functional Ecology 557–562 (2002).

72. H. T. Pinheiro, et al., Island biogeography of marine organisms. Nature 549, 82–85 (2017).

73. D. R. Bellwood, T. P. Hughes, Regional-scale assembly rules and biodiversity of coral reefs. Science 292, 1532–1535 (2001).

74. C. Chaudhary, H. Saeedi, M. J. Costello, Bimodality of latitudinal gradients in marine species richness. Trends in Ecology & Evolution 31, 670–676 (2016).

75. M. Messié, et al., Potential new production estimates in four eastern boundary upwelling ecosystems. Progress in Oceanography 83, 151–158 (2009).

76. E. Puccinelli, et al., Omega-3 pathways in upwelling systems: the link to nitrogen supply. Frontiers in Marine Science 8, 664601 (2021).

77. F. P. Chavez, M. Messié, A comparison of eastern boundary upwelling ecosystems. Progress in Oceanography 83, 80–96 (2009).

78. J. A. Santora, et al., Persistence of trophic hotspots and relation to human impacts within an upwelling marine ecosystem. Ecological Applications 27, 560–574 (2017).

79. M. A. Rex, Community structure in the deep-sea benthos. Annual Review of Ecology and Systematics 12, 331–353 (1981).

80. J. M. Chase, M. A. Leibold, Spatial scale dictates the productivity–biodiversity relationship. Nature 416, 427–430 (2002).

81. E. J. Sayer, L. M. Sutcliffe, R. I. Ross, E. V. Tanner, Arthropod abundance and diversity in a lowland tropical forest floor in Panama: the role of habitat space vs. nutrient concentrations. Biotropica 42, 194–200 (2010).

82. M. Kaspari, S. P. Yanoviak, Biogeochemistry and the structure of tropical brown food webs. Ecology 90, 3342–3351 (2009).

83. W. S. Harpole, et al., Addition of multiple limiting resources reduces grassland diversity. Nature 537, 93–96 (2016).

84. S. J. Brandl, et al., Demographic dynamics of the smallest marine vertebrates fuel coral reef ecosystem functioning. Science 364, 1189–1192 (2019).

85. M. Kaspari, et al., Multiple nutrients limit litterfall and decomposition in a tropical forest. Ecology letters 11, 35–43 (2008).

86. S. Hättenschwiler, H. B. Jørgensen, Carbon quality rather than stoichiometry controls litter decomposition in a tropical rain forest. Journal of Ecology 98, 754–763 (2010).

87. O. P. L. Vindstad, T. Birkemoe, R. A. Ims, A. Sverdrup-Thygeson, Environmental conditions alter successional trajectories on an ephemeral resource: a field experiment with beetles in dead wood. Oecologia 194, 205–219 (2020).

88. A. C. Cleary, E. G. Durbin, T. A. Rynearson, J. Bailey, Feeding by Pseudocalanus copepods in the Bering Sea: trophic linkages and a potential mechanism of niche partitioning. Deep Sea Research Part II: Topical Studies in Oceanography 134, 181–189 (2016).

89. X. He, M. Lei, F. Cheng, S. Hu, In situ food compositions reveal niche partitioning in small marine cladocerans and copepods in Daya Bay, South China Sea. Mar. Ecol. Prog. Ser. 716, 47–61 (2023).

90. A. Novotny, S. Zamora-Terol, M. Winder, DNA metabarcoding reveals trophic niche diversity of micro and mesozooplankton species. Proceedings of the Royal Society B 288, 20210908 (2021).

91. J. W. Dacey, S. G. Wakeham, Oceanic dimethylsulfide: production during zooplankton grazing on phytoplankton. Science 233, 1314–1316 (1986).

92. J. L. DeBose, G. A. Nevitt, A. H. Dittman, Rapid communication: experimental evidence that juvenile pelagic jacks (Carangidae) respond behaviorally to DMSP. Journal of chemical ecology 36, 326–328 (2010).

93. S. Kowalewsky, M. Dambach, B. Mauck, G. Dehnhardt, High olfactory sensitivity for dimethyl sulphide in harbour seals. Biology letters 2, 106–109 (2006).

94. G. A. Nevitt, M. Losekoot, H. Weimerskirch, Evidence for olfactory search in wandering albatross, Diomedea exulans. Proceedings of the National Academy of Sciences 105, 4576–4581 (2008).

95. J. M. Casey, et al., DNA metabarcoding marker choice skews perception of marine eukaryotic biodiversity. Environmental DNA 3, 1229–1246 (2021).

96. S. Piao, et al., Changes in climate and land use have a larger direct impact than rising CO2 on global river runoff trends. Proceedings of the National academy of Sciences 104, 15242–15247 (2007).

97. P. Souza, S. Brandl, Historic Freshwater Inflow Silences an Estuarine Ecosystem. Estuaries and Coasts 48, 1–15 (2025).

98. A. Liess, et al., Terrestrial runoff boosts phytoplankton in a Mediterranean coastal lagoon, but these effects do not propagate to higher trophic levels. Hydrobiologia 766, 275–291 (2016).

99. O. F. Rowe, et al., Climate change–induced terrestrial matter runoff may decrease food web production in coastal ecosystems. Limnology and Oceanography (2025).

100. E. B. Barbier, et al., The value of estuarine and coastal ecosystem services. Ecological monographs 81, 169–193 (2011).

101. M. Leray, et al., A new versatile primer set targeting a short fragment of the mitochondrial COI region for metabarcoding metazoan diversity: application for characterizing coral reef fish gut contents. Frontiers in zoology 10, 34 (2013).

102. J. Geller, C. Meyer, M. Parker, H. Hawk, Redesign of PCR primers for mitochondrial cytochrome c oxidase subunit I for marine invertebrates and application in all-taxa biotic surveys. Molecular ecology resources 13, 851–861 (2013).

103. E. Bolyen, et al., Reproducible, interactive, scalable and extensible microbiome data science using QIIME 2. Nature biotechnology 37, 852–857 (2019).

104. M. Martin, Cutadapt removes adapter sequences from high-throughput sequencing reads. *EMBnet*. journal 17, 10–12 (2011).

105. B. J. Callahan, et al., DADA2: High-resolution sample inference from Illumina amplicon data. Nature methods 13, 581–583 (2016).

106. S. F. Altschul, W. Gish, W. Miller, E. W. Myers, D. J. Lipman, Basic local alignment search tool. Journal of molecular biology 215, 403–410 (1990).

107. M. Leray, N. Knowlton, R. J. Machida, MIDORI2: A collection of quality controlled, preformatted, and regularly updated reference databases for taxonomic assignment of eukaryotic mitochondrial sequences. Environmental Dna 4, 894–907 (2022).

108. S. Ratnasingham, P. D. Hebert, BOLD: The Barcode of Life Data System (http://www.barcodinglife.org). Molecular ecology notes 7, 355–364 (2007).

109. P. Pappalardo, et al., Taxon-specific BLAST percent identity thresholds for identification of unknown sequences using metabarcoding. Methods in Ecology and Evolution 16, 2380–2394 (2025).

110. J. Xue, C. Lee, S. G. Wakeham, R. A. Armstrong, Using principal components analysis (PCA) with cluster analysis to study the organic geochemistry of sinking particles in the ocean. Organic Geochemistry 42, 356–367 (2011).

111. F. Keck, M. Couton, F. Altermatt, Navigating the seven challenges of taxonomic reference databases in metabarcoding analyses. Molecular Ecology Resources 23, 742–755 (2023).

112. W. Viechtbauer, Conducting meta-analyses in R with the metafor package. Journal of statistical software 36, 1–48 (2010).

113. P.-C. Bürkner, brms: An R package for Bayesian multilevel models using Stan. Journal of statistical software 80, 1–28 (2017).

114. J. E. Byrnes, L. E. Dee, Causal inference with observational data and unobserved confounding variables. Ecology Letters 28, e70023 (2025).

115. R. McElreath, Statistical rethinking: A Bayesian course with examples in R and Stan (Chapman and Hall/CRC, 2018).

116. R Core Team, R: A language and environment for statistical computing. R Foundation for Statistical Computing, Vienna, Austria. URL https://www.R-project.org/. (2022). Deposited 2022.

117. J. Oksanen, et al., The vegan package. Community ecology package 10 (2007).

118. H. Wickham, et al., Welcome to the Tidyverse. Journal of Open Source Software 4, 1686 (2019).

