## Supplemental Material for "Energy quality shapes biodiversity across coastal oceans"

### Electronic Supplementary Material

**This PDF file includes:**

Supporting Methods – Site Descriptions
Supporting Methods – Supplemental Analyses
Figures S1 to S3
Table S1

#### Supplemental Methods

##### *Site descriptions*

Below, we provide detailed descriptions of all sampling locations and their sites.

*Texas Gulf Coast, Texas, USA*: Sampling was conducted in the Mission-Aransas National Estuarine Research Reserve (MANERR). The five sampling sites corresponded to the NERR's system-wide monitoring platforms (SWMP stations: [27.9746º, -97.0239º], [28.08403º, -97.2009º], [28.13234, -97.03444], [28.1385, -96.82847], [27.838, -97.05026]), spanning an estuarine-marine gradient from the western reaches of Copano Bay through Aransas and Mesquite Bay to the Port Aransas Ship Channel, which connects the vast estuarine landscape to the Gulf of Mexico. The Mission-Aransas estuary is a shallow, largely wind-driven system of open bays that is fed by the Mission and Aransas Rivers, and it encompasses a wide range of estuarine habitats, such as oyster reefs, seagrass meadows, and marshes. Copano Bay, which is closest to the river inlets, is generally the least saline part of the estuary, while Mesquite and Aransas Bay have higher salinities due to increased exchange with the open Gulf. The site within the ship channel is highly marine. Sampling in South Texas was performed on June 29, 2023.

*Gulf of Panama, Panama*: The Gulf of Panama is part of the Panama Bight and is characterized by strong seasonal oceanographic variability. During the dry season (January–April), northerly trade winds drive recurrent upwelling brings cooler, nutrient-rich waters to the surface, contrasting with the warmer, more oligotrophic conditions typical of the wet season (May–December). The broader coastal zone of the Gulf of Panama includes extensive mangrove systems that receive high sediment and nutrient inputs from surrounding watersheds and experience strong tidal amplitudes. Sampling was conducted in July 2023 along a coastal–offshore transect in the central Gulf of Panama extending from east of Flamenco Island to waters north of Otoque Island. The five sampling sites (from north to south) were Flamenco (8.9123º, −79.4998º), Balboa (8.8493º, −79.5236º), Taboga (8.7959º, −79.5343º), Chama (8.7006º, −79.5491º), and Otoque (8.6523º, −79.5694º). These stations are located at increasing distance from the heavily trafficked Pacific entrance of the Panama Canal, spanning a gradient from urban-influenced coastal waters to more offshore island environments. The northernmost sites near Flamenco and Balboa are influenced by intense shipping activity associated with canal operations and runoff from the Panama City metropolitan region. Moving offshore toward Taboga, Chama, and Otoque, the sites transition to more oceanic conditions around islands that support sandy beaches, soft-sediment bottoms, rocky substrates, and local coral communities.

*Bocas del Toro, Panama:* Bahía Almirante is a multiple inlet bay on the Caribbean coast of Panama. Consistently warm, the region experiences two wet and two dry seasons per year, with as much as 3m of rainfall. However, freshwater inputs from direct rainfall and rivers are roughly equal, with rivers from small catchments emptying into the bay. Occasionally the river plume of the adjacent Rio Changuinola is pushed into the bay through the Boca del Drago Channel. The bay experiences seasonal hypoxia at depth driven by stratification and infrequent bottom-water renewal. The bay is boarded by red mangroves and has extensive seagrass beds, some of which overlie extensive peat deposits. The sampling depth was 20-25m at all five sites. Sites were situated along the Bahia Almirante coast of Isla Colon (sites 1-3) adjacent to the mangrove fringe. The most highly impacted site also had the greatest oceanic exchange on the outer side of Isla Colon adjacent to Bocas del Toro town (site 4). The site with least oceanic influence was well inside Bahia Almirante at Punta Juan on San Cristobal Island (site 5). The site coordinates were:

(1) 09.36367º, -82.28458º; (2) 09.34824º, -82.26259º; (3) 09.32973º, -82.25446º; (4) 09.34379º, -82.24229º; (5) 09.30128º, -82.29413º.

*Indian River Lagoon, Florida, USA*: The Indian River Lagoon (IRL) is a shallow, microtidal estuary located along Florida's Atlantic coast and is characterized by strong gradients in salinity, water residence time, and urbanization. Sampling was conducted in the central IRL near the Ft. Pierce Inlet. Five sampling sites were selected along a north-south transect spanning an inlet-to-lagoon gradient: (1) 27.4695º, -80.3039º, (2) 27.4878º, -80.3050º, (3) 27.5368º, -80.3487º, -27.5603º,(4) -80.3393º, and (5) 27.4423º, -80.2978º. Sites 3, 4, and 5 coincide with long-term MarineGEO seagrass monitoring locations and represent relatively less urbanized regions of the lagoon with established submerged aquatic vegetation. Site 2 is situated within a heavily urbanized portion of the IRL, while Site 1 lies closest to the Ft. Pierce Inlet and is most strongly influenced by marine exchange. Together, these sites span gradients in hydrology, land use, marine influence. The sites were selected to provide gradient in water-column productivity and biodiversity.

*Madeira Archipelago, Portugal.* The Madeira Archipelago is located in the Atlantic Ocean at the dynamic boundary of the North Atlantic Subtropical Gyre where the northward-flowing Azores Current intersects with the eastward-flowing Canary Current. It lies approximately 1,000 km southwest of mainland Portugal and forms part of the Macaronesian biogeographic region, together with the Azores, Canary Islands, and Cape Verde archipelagos. Sampling was conducted at five stations spanning gradients of demographic pressure, hydrology, land use, and marine influence. Four stations were located along the southern coast of Madeira Island, from east to west: Caniçal (32.741266º, -16.709676º), Funchal (32.645703º, -16.911222º), Ribeira Brava (32.668044º, -17.063515º), and Paúl do Mar (32.751253º, -17.224215º). One additional station was located on the northwestern coast at Porto Moniz (32.866133º, -17.164933º). Funchal, the island's capital, exhibits the highest levels of human population density and anthropogenic activity, followed by Ribeira Brava. Caniçal, Paul do Mar, and Porto Moniz represent sites with comparatively lower population density and human activity, with the exception of Caniçal, which hosts a commercial harbor.

*British Columbia, Canada*: Sampling was conducted at five sites around Quadra Island, one of the largest islands in the northern end of the Salish Sea off the east coast of Vancouver Island. Quadra's shoreline is largely bedrock and cobble, and high tidal exchanges combined with narrow passages create strong tidal currents along much of the northern and western shores of the island. The southern and eastern shores are less exposed to strong tidal currents, but they are fully exposed to the Salish Sea in the south. The oceanographic conditions around Quadra Island are mostly influenced by water coming from the Juan de Fuca Strait, located > 300 km to the south and moving northward through the Strait of Georgia. There is considerable rainfall during the winter months, contrasted with periods of dryer, warmer weather in the summer months. Most of the freshwater outputs from Quadra Island are small creeks or rivers, though there can be considerable seasonal freshwater fluxes from the surrounding glacier-fed watersheds on the mainland and Vancouver Island. Sampling sites were selected based on shore access via docks but also cover a wide spatial gradient (~22 km north to south, ~15 km east-west), as well as a range of human impacts. The five sites were located at: (1) Heriot Bay [50.10311º, -125.21133º], a marina, ferry terminal and small human establishment on the east side of the island, (2) April Point [50.06256º, -125.22728º], a small marina on the west side of the island in a sheltered bay, (3) Quathiaski Cove [50.04252º, -125.21570º], a marina, ferry terminal, and small human establishment on the west, (4) Surge Narrows [50.213194º, -125.147790º], a small boat launch in the northeast subject to low human impact and high current flows, and (5) Granite Bay

[50.237746º, -125.307184º], a small dock and boat launch in the north subject to low human impact and in a large, sheltered bay.

*Kāne'ohe, O'ahu, Hawai'i*: Sampling was conducted at five sites in Kāne'ohe on the windward side of the island of O'ahu, Hawai'i, on June 21, 2023. The five sampling sites are part of two time-series programs. The most nearshore site (site 1, code "KK", 21.43582ºN, -157.80524ºE) is part of the He'eia National Estuarine Research Reserve's System-Wide Monitoring Program (SWMP). The four other sites (site 2, code "HP1", 21.43688º, -157.80331º; site 3, code "SR8", 21.44983333º, -157.79322º; site 4, code "SR2", 21.4691667º, -157.77752º; site 5, code "STO1", 21.4829º, -157.7663º) are part of the Kāne'ohe Bay Time-series (KByT). SWMP data is publicly available and can be found at the Centralized Data Management Office website (https://cdmo.baruch.sc.edu/). These sites span nearshore, estuarine sites immediately adjacent to an ancient Hawaiian fishpond (KK and HP1), outward in a transect in Kāne'ohe Bay (SR8, SR2), and toward the oligotrophic North Pacific Subtropical Gyre (STO1). Kāne'ohe Bay is a subtropical, semi-enclosed embayment representing a sharp gradient of waters influenced by nearshore, land-based impacts, to clear, low-nutrient waters of the Pacific Ocean.

*East Tasmania, Australia:* Sampling was conducted along the east coast of Tasmania across five coastal sites spanning exposed open-coast and more sheltered nearshore environments. The sampling sites were Bolton Beach (42.953239°, 147.355381°), Triabunna (42.523206°, 147.912275°), Shelly Beach (41.869689°, 148.302850°), Spring Beach (41.248431°, 148.331069°), and Rheban Beach (43.283939°, 147.166669°).Together, these sites span a latitudinal gradient along Tasmania's east coast and include open sandy beaches directly exposed to Tasman Sea conditions as well as sheltered waters within coastal embayments, including Mercury Passage, which separates mainland Tasmania from Maria Island. The region is characterized by cool-temperate coastal conditions and is influenced by the southward extension of the East Australian Current, which intermittently introduces warmer oceanic water along the coast.

*Chesapeake Bay, Maryland, USA*: Sampling was conducted in the Rhode River, Maryland. The Rhode River is a shallow sub-embayment of the Chesapeake Bay that is fed by several freshwater creeks and encompasses a range of estuarine habitats including marshes, submerged aquatic vegetation, oysters, and sandy and mud bottoms. Located approximately two thirds of the way up the bay and 200 km from the ocean in the mesohaline region, salinity is largely driven by bay-wide conditions, which vary seasonally and annually with the amount of freshwater inflow from the Susquehanna River, as well as the local watershed. Sampling sites were selected along a gradient of the river toward the bay, ending at the mouth of the river. The five sampling sites were Fox Point (38.8819º, -76.5443º), SERC Dock (38.8854º, -76.5415º), Big Island (38.8838º, -76.5364º), Canninghouse Bay (38.8758º, -76.5263º), and Mouth Mud (38.8660º, -76.5213º). Sampling was performed on June 21, 2023.

*Hong-Kong*: Sampling was conducted in Tolo Harbor, Hong Kong. The sampling sites were Central Island (22.437953°, 114.222564°), Che Lei Pai (22.462097°, 114.290428°), Luk Wu Tung (22.487953°, 114.311731°), Port Island (22.501383°, 114.355992°) and Tung Ping Chau (22.550931°, 114.430953°). Tolo harbor is an enclosed bay in northeast Hong Kong that was historically a nursery ground for commercially important fishes, but it has lost foundational species due to anthropogenic pressure from coastal development and nutrient pollution. These sites span a gradient of water quality and habitat degradation, from highly impacted inner harbor locations with low salinity and high terrestrial organic matter input to more oceanic, higher salinity sites with greater coral cover and marine influence. Sampling was performed on May 30, 2023.

*Central Peruvian Coast, Peru*: Sampling was conducted in subtidal waters off Pampa Larga, a ~16 km sandy shoreline located in Cañete, Lima, Peru, within the influence of a liquefied natural gas (LNG) port terminal. Sampling site coordinates were: Site 1: 77.400508°, −13.261507°; Site 2: 80.242873°, −13.192535°; Site 3: 77.905701°, −13.224319°; Site 4: 75.936435°, −13.311807°; and Site 5: 81.213043°, −13.289160°. This coastal system lies within the Humboldt Current, a highly productive eastern boundary upwelling system. Submerged habitats are dominated by extensive soft-bottom sediment flats interspersed with occasional rocky reefs, as well as artificial hard substrates associated with port infrastructure (e.g., jetty pilings and two breakwaters). The shallow zone (0–8 m) is primarily composed of sand and gravel, transitioning to finer sediments (silt and mud) at depths of 8–15 m. Local oceanographic conditions are driven by seasonal upwelling, wave action, and coastal currents, which promote vertical mixing and sediment resuspension. Waters are typically cold, saline, and nutrient-rich, with moderate to high turbidity. The region is arid, with minimal freshwater input, although episodic climatic variability (e.g., El Niño events) can influence temperature and productivity. Sampling sites were distributed across the terminal's area of influence and adjacent control zones.

*Lower Chesapeake, Virginia, USA:* Sampling was conducted in shallow subtidal waters of Mobjack Bay and the York River in the lower Chesapeake Bay, an estuarine drowned river mouth ecosystem. Mobjack Bay is a tidal embayment within the Chesapeake, and samples were collected near Guinea Marsh (37.285212º, -76.350734º), New Point Comfort (37.30025º, -76.278916º), and Four Point Marsh (37.338405º, -76.398098º). To the south of Mobjack Bay, the York River feeds into the Chesapeake and samples were collected near Perrin Creek (37.262851º, -76.409104º) and south of Goodwin Island (37.204631º, -76.398121º). The region is characterized by spartina marshes (*Spartina alterniflora*), fringing and submerged oyster reefs (*Crassostrea virginica*), and seagrass meadows composed of eelgrass (*Zostera marina*) and widgeongrass (*Ruppia maritima*). The system is polyhaline, with a salinity of 18-22 PSU, and is typically characterized by eutrophic conditions. All samples were collected at a depth of approximately 2m on June 28, 2023.

##### *Supplemental analyses*

We ran several additional analyses to examine the robustness of our findings. First, we re-ran our global and regional regression analyses for all taxa using only locations (global model) and sites (regional model) that displayed average salinity values >30. To do so, we filtered each dataset and ran the same model that was previously specified (see *Statistical Analysis* section in the *Methods*). The resulting datasets had fewer replicates (N = 9 for the global analysis, N = 46 for the regional analysis), but revealed the same effects as the full model (see *Results*). This reveals that the obtained patterns are not simply due to a dichotomy between select estuarine locations and the remaining study sites.

Furthermore, we tested whether univariate variables describing the quantity and quality of energy had divergent or similar effects on. biodiversity as the latent variables we derived from the PCA. Due to issues with collinearity (as predictable based on the PCA), we tested carbon concentrations (μmol C $L^{-1}$) and nitrogen concentrations (μmol N $L^{-1}$) as raw descriptors of energy quantity in separate models, and we tested C:N ratio and the degradation index as variables describing energy quality separately, each at global and regional spatial scales. Carbon and nitrogen concentrations both showed fully comparable, analogous results to the latent energy quantity variable at the global and regional scale: both showed clear, negative correlations (C global = -0.76 [-0.83, -0.69]; C regional = -0.38 [-0.73, -0.01]; N global = -0.73 [-0.80, -0.66]; N regional = -0.55 [-0.89, -0.16]) with biodiversity across scales as expected given their extremely high collinearity and loading on PC1. For C:N ratio and $\delta^{13}C$, results differed between regional and global models. C:N ratio (0.23 [0.09, 0.36]) and the degradation index (-0.30 [-0.44, -0.16]) both showed clear relationships with biodiversity at the global scale, but no clear relationships at the regional scale (C:N ration: -0.08 [-0.24, 0.10]; degradation index: -0.02 [-0.28, 0.26]). Thus, while the energy quantity index is directly comparable to the raw variables that define it, the latent energy quality variable is useful in capturing multiple facet of resource accessibility and nutritional value that are better captured as a multivariate index.

#### Figures

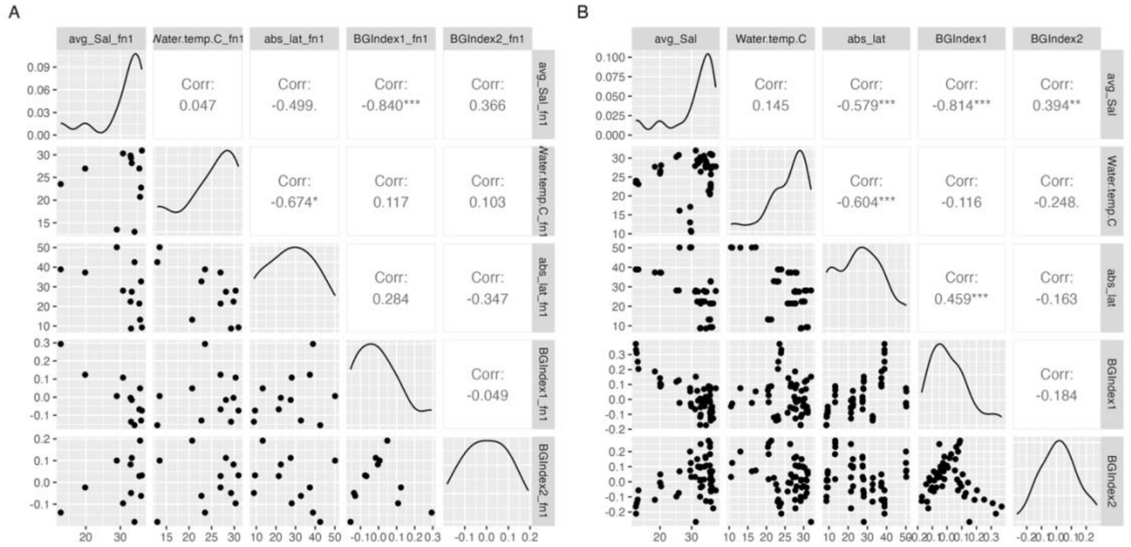


**Figure S1:** Correlation coefficients for the explanatory variables used in the analyses at the location (A) and site level (B). Due to high correlation coefficients for temperature (Water.temp.C) and absolute latitude (abs_lat) (-0.674 and -0.604, respectively), salinity (avg_Sal) and energy quantity (BGIndex1) (-0.840 and -0.814, respectively), we dropped absolute latitude from all models and evaluated salinity in a separate model.

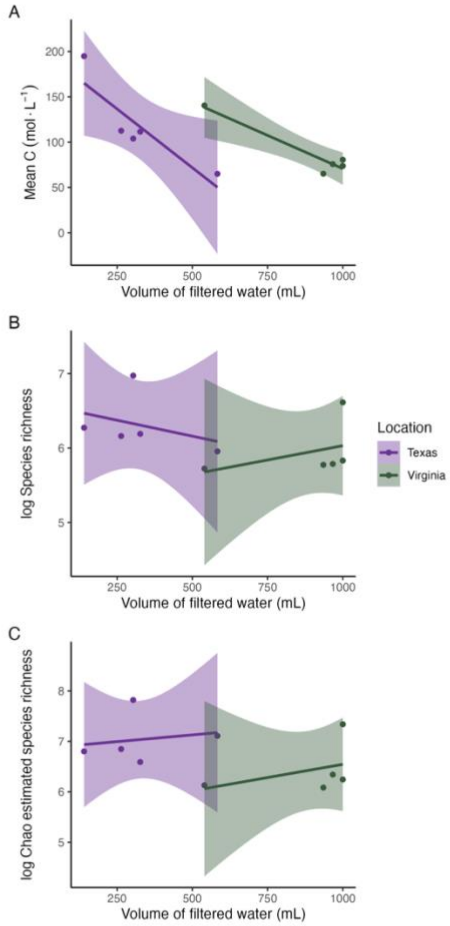


**Figure S2:** Relationship between the volume of filtered water (x-axes) and the total organic carbon concentration (A), amplicon sequence variant (ASV) richness (proxy for species richness) (B), and the incidence-based rarefaction results for the ASVs (C) in two locations (Texas and Virginia) where we sampled different volumes of water. Despite a weak negative relationship between carbon concentration and filter volume (given the likelihood of organic carbon clogging filter pores over time), no relationship is apparent between water volume and biodiversity.

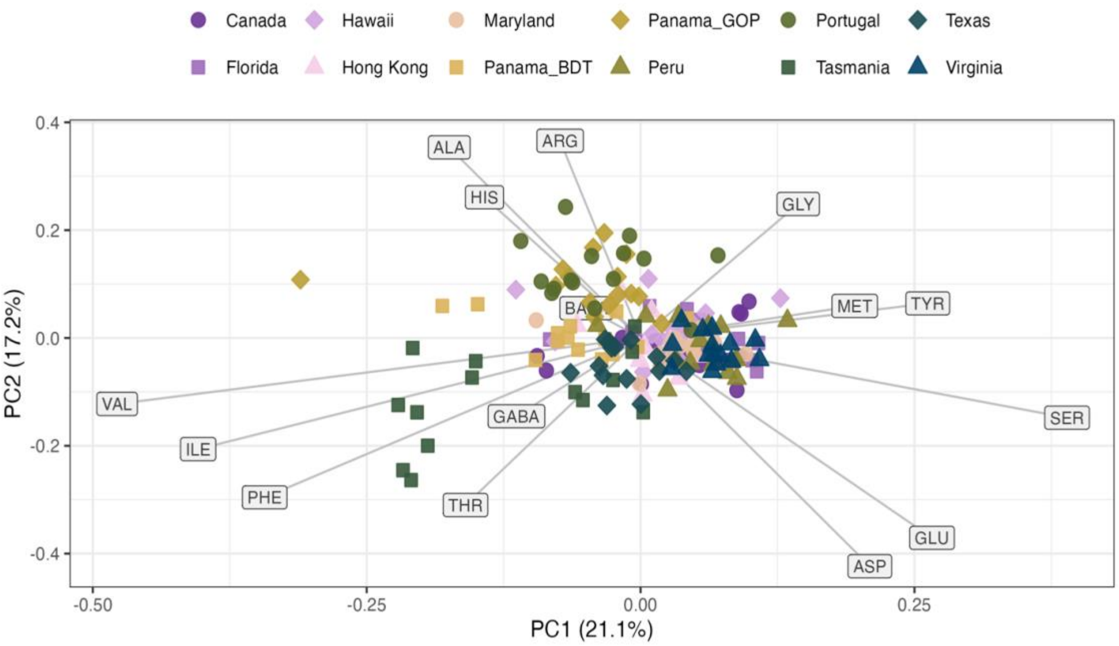


**Figure S3:** Principal component analysis (PCA) delineating the degradation index of organic matter based on the arrangement of indicator amino acids in bivariate space. The he primary axis of the degradation index is defined by the arrangement of BALA/GABA and ASP/GLU as the main indicators of organic matter lability. Panama_GOP = Gulf of Panamá, Panamá, Panama_BDT = Bocas del Toro, Panamá. Amino acids are as follows: aspartic acid (ASP), glutamic acid (GLU), histidine (HIS), serine (SER), arginine (ARG), glycine (GLY), threonine (THR), β-alanine (BALA), alanine (ALA), tyrosine (TYR), γ-aminobutyric acid (GABA), methionine (MET), valine (VAL), phenylalanine (PHE), isoleucine (ILE), and leucine (LEU).

**Table S1:** Classification of taxa in the combined reference library to remove predominantly terrestrial classes. Only taxa confidently assigned to the "Terrestrial" category were removed. "Mixed" indicate taxa that are both terrestrial and marine, or that little is known about the habitat of the taxon. The *Count* column indicates the prevalence of each taxon in the combined reference database.

| ***Class*** | ***Count*** | ***Ecology*** |
|---|---|---|
| Insecta | 7278777 | Terrestrial |
| Arachnida | 388990 | Terrestrial |
| Actinopterygii | 318438 | Marine |
| Collembola | 186478 | Terrestrial |
| Malacostraca | 171201 | Marine |
| Gastropoda | 165218 | Marine |
| Mammalia | 130284 | Mixed |
| Bivalvia | 56662 | Marine |
| Aves | 50506 | Mixed |
| Clitellata | 50035 | Mixed |
| Polychaeta | 35899 | Marine |
| Amphibia | 33491 | Terrestrial |

| | | |
|---|---|---|
| Copepoda | 26702 | Marine |
| Reptilia | 23827 | Terrestrial |
| Elasmobranchii | 23372 | Marine |
| Florideophyceae | 22799 | Marine |
| Thecostraca | 20254 | Marine |
| Chromadorea | 19072 | Marine |
| Branchiopoda | 17847 | Marine |
| Anthozoa | 13302 | Marine |
| Ophiuroidea | 12829 | Marine |
| Cephalopoda | 11811 | Marine |
| Trematoda | 10592 | Marine |
| Asteroidea | 9986 | Marine |
| Demospongiae | 6728 | Marine |
| Ascidiacea | 6595 | Marine |
| Diplopoda | 6497 | Terrestrial |
| Hydrozoa | 6479 | Marine |
| Monogononta | 5951 | Mixed |
| Ostracoda | 5040 | Marine |
| Echinoidea | 4993 | Marine |
| Cestoda | 4876 | Mixed |
| Scyphozoa | 4829 | Marine |
| Chilopoda | 4498 | Terrestrial |
| Crinoidea | 4449 | Marine |
| Bdelloidea | 4313 | Mixed |
| Holothuroidea | 4309 | Marine |
| Polyplacophora | 4270 | Marine |
| Pycnogonida | 3627 | Marine |
| Oomycota | 3424 | Mixed |
| Peronosporales | 3246 | Mixed |
| Hoplonemertea | 2761 | Mixed |
| Gymnolaemata | 2648 | Mixed |
| Palaeacanthocephala | 2422 | Marine |
| Aconoidasida | 2418 | Mixed |
| Rhabditophora | 2407 | Mixed |
| Pilidiophora | 2272 | Mixed |
| Eutardigrada | 1771 | Mixed |
| Udeonychophora | 1717 | Terrestrial |
| Pythiales | 1510 | Mixed |
| Sagittoidea | 1481 | Marine |
| Dinophyceae | 1257 | Mixed |

| | | |
|---|---|---|
| Testudines | 1017 | Mixed |
| Enoplea | 1007 | Mixed |
| Bangiophyceae | 1005 | Mixed |
| Monogenea | 927 | Mixed |
| Sipuncula | 899 | Marine |
| Bacillariophyceae | 856 | Mixed |
| Heterotardigrada | 797 | Mixed |
| Conoidasida | 716 | Mixed |
| Petromyzonti | 662 | Marine |
| Palaeonemertea | 562 | Mixed |
| Oligohymenophorea | 552 | Mixed |
| Eurotiomycetes | 552 | Terrestrial |
| Holocephali | 525 | Marine |
| Eucycliophora | 472 | Mixed |
| Gordioida | 447 | Mixed |
| Leptocardii | 436 | Marine |
| Symphyla | 411 | Terrestrial |
| Myxini | 388 | Marine |
| Gastrotricha_class_incertae_sedis | 375 | Mixed |
| Cyclorhagida | 375 | Mixed |
| Eoacanthocephala | 368 | Mixed |
| Crocodylia | 329 | Mixed |
| Tentaculata | 321 | Mixed |
| Acoelomorpha | 321 | Mixed |
| Sordariomycetes | 312 | Terrestrial |
| Protura | 292 | Terrestrial |
| Agaricomycetes | 285 | Terrestrial |
| Diplura | 259 | Terrestrial |
| Ichthyostraca | 227 | Marine |
| Sarcopterygii | 217 | Marine |
| Cubozoa | 209 | Marine |
| Saccharomycetes | 209 | Terrestrial |
| Merostomata | 203 | Mixed |
| Leotiomycetes | 202 | Terrestrial |
| Solenogastres | 191 | Mixed |
| Enteropneusta | 191 | Mixed |
| Homoscleromorpha | 176 | Mixed |
| Acoela | 169 | Mixed |
| Saprolegniales | 165 | Mixed |
| Scaphopoda | 150 | Mixed |

| | | |
|---|---|---|
| Priapulimorpha | 150 | Mixed |
| Macrodasyida | 148 | Mixed |
| Hexactinellida | 137 | Mixed |
| Lingulata | 136 | Mixed |
| Archiacanthocephala | 133 | Mixed |
| Thaliacea | 128 | Marine |
| Pauropoda | 120 | Terrestrial |
| Nuda | 119 | Mixed |
| Priapulida_class_incertae_sedis | 113 | Mixed |
| Rhynchonellata | 103 | Mixed |
